# Dengue virus NS1 undergoes partial nuclear translocation to modulate host transcription and support viral replication

**DOI:** 10.64898/2026.04.13.718202

**Authors:** César Pacheco, Raymundo Cruz, Christopher D. Wood, Eva Zusinaite, Andres Merits, Rodolfo Gamaliel Ávila Bonilla, Refugio García-Villegas, Juan E. Ludert

## Abstract

The dengue virus (DENV) non-structural protein 1 (NS1) is a glycoprotein highly conserved among mosquito-borne orthoflaviviruses. NS1 is typically localized in the lumen of the endoplasmic reticulum, where it forms part of the replication complexes, and is also exposed at the plasma membrane. In addition, NS1 is secreted as a lipoprotein. Here, using a combination of approaches, including confocal microscopy with deconvolution, in situ analysis, and biochemical cell fractionation, we show that a substantial fraction of NS1 (up to 30%) translocates to the nucleus during infection. We identified a conserved, structurally exposed bipartite nuclear localization signal (NLS) within NS1. Pharmacological inhibition with ivermectin and site-directed mutagenesis of the NLS in recombinant confirmed that nuclear import of NS1 is an active process, dependent on the classical importin α/β pathway. Notably, both dimeric and multimeric forms of NS1 were detected in the nucleus in association with nuclear lamin. Introduction of the NLS mutations into DENV2 infectious clones resulted in a non-viable virus. Production of virus progeny and completion of the replicative cycle by the mutant genomes could be rescued by trans-complementation with wild-type NS1, but not with an NLS-mutated NS1, indicating that an NS1 nuclear phase is required for a productive infection. Transcriptomic analysis by RNA-seq further revealed that NS1 functions depend on its subcellular location. Nuclear NS1 induced the overexpression of genes associated with DNA-binding transcription factors, whereas NLS-mutated NS1, retained in the cytoplasm, failed to induce these genes and instead triggered pro-inflammatory and metabolic responses. Together, these findings reveal a previously unrecognized nuclear phase of NS1 that is required for an efficient viral life cycle, redefining NS1 as a modulator of the host transcriptional environment. These findings also suggest new avenues for antiviral and vaccine development.

**Authors summary:** Dengue virus NS1 is a glycoprotein of approximately 45-50 kDa that rapidly dimerizes after proteolytic maturation. Dimeric NS1 is located in the lumen of the endoplasmic reticulum where it acts as a scaffold component of the viral replication complexes. In addition, NS1 is secreted from infected cells as a tetramer or hexamer and circulates in the serum of infected individuals during the acute phase of dengue disease. Circulating NS1 is widely used as a diagnostic marker and has also been associated with dengue pathogenesis through several mechanisms. Here, we expand the current understanding of DENV NS1 by identifying a previously unrecognized and essential nuclear location of this protein. We show that NS1 contains a conserved nuclear localization signal that mediates import into the nucleus via the classical import pathway. Using wild-type and NLS-mutated infectious clones, we demonstrated that nuclear localization of NS1 is required for completion of the DENV replicative cycle. Transcriptomic analysis further revealed that nuclear NS1 promotes the expression of host genes involved in nucleic acid metabolism, whereas retention of NS1 in the cytoplasm triggers an antiviral and inflammatory response. Together, these findings identify the nucleus as an important site of dengue virus-host interactions and redefine NS1 as a regulator of the host transcriptional environment during infection.

## Introduction

Dengue virus (DENV, *Orthoflavivirus denguei*) is a mosquito-borne flavivirus that represents a significant threat to global health. In its urban cycle, DENV is transmitted mainly by *Aedes aegypti* and *Aedes albopictus* mosquitoes. Dengue disease presents with a wide spectrum of clinical symptoms, ranging from asymptomatic infections, to dengue fever without or with alarm symptoms, to severe dengue, which can be potentially fatal (1). The DENV genome is composed of a monocistronic RNA of positive polarity, of approximately 10Kb, which encodes a polyprotein that is processed co– and post transcriptionally into three structural proteins (C, prM, E) that conform the virion, and seven nonstructural proteins (NS1, NS2A, NS2B, NS3, NS4A, NS4B, NS5). The nonstructural proteins are critical for orchestrating viral replication, virion assembly, and antagonizing the cell’s innate immune response (2).

The nonstructural protein 1 (NS1) is a glycoprotein of approximately 45-50 kDa, highly conserved across mosquito-borne orthoflaviviruses, with multiple roles during viral infection. During synthesis, NS1 is translocated into the lumen of the endoplasmic reticulum (ER), where it rapidly forms membrane-associated dimers that serve as scaffolds for replication complexes (3). A role for intraluminal NS1 in virion morphogenesis has also been proposed (4,5) NS1 is also trafficked to the plasma membrane by an unknown route, and remains attached to the outer leaflet of the plasma membrane via a glycosylphosphatidylinositol (GPI) tail (6). Recently, NS1 was observed in the cell cytoplasm of infected mosquito cells, interacting with lipid droplets (7). Finally, NS1 is secreted from infected cells as tetramers and hexamers associated with lipids, making soluble NS1 a true lipoprotein (8,9). Soluble NS1 circulates in the serum of patients at high concentrations during the acute phase of the disease, making it a robust diagnostic marker (10). However, circulating NS1 has been implicated in dengue pathogenesis by different mechanisms that include activation of proinflammatory responses via TLR4 activation (11), disruption of the complement system (12), induction of endothelial hyperpermeability (13), immune dysregulation and facilitation of viral replication (14,15), and even the generation of self-reactive antibodies (13,16,17).

Although not fully understood, many of the properties and multiple functions of DENV NS1 in the ER, the plasma membrane, and in the extracellular space have been studied and characterized (18–21). However, there is evidence suggesting that NS1 may have additional subcellular locations in both infected mosquito and mammalian cells. For example, immune electron microscopy studies have described the presence of DENV NS1 inside cell nuclei (22,23). Biochemical cell fractionation of DENV-infected human endothelial cells showed NS1 to be present in membrane/organelle, nuclear, and cytoskeletal fractions (24). In agreement, recent protein-protein interactome maps obtained in mammalian and mosquito cells identified several components of the nuclear import and export machinery, such as karyopherins and nucleoporins, as well as several proteins of the nuclear pore complex as part of the NS1 interactome (25,26). Despite this compelling evidence, the dynamics, import route, functional relevance, and consequences for the DENV life cycle of the association of NS1 with the nucleus remain totally unexplored.

In this study, we examine the presence, traffic route, and biological functions of DENV2 NS1 within the nucleus of infected or transiently transfected mammalian cells. The results show that NS1 accumulates in the nucleus early after infection, accounting for up to 30% of the total intracellular NS1 pool, using a functional and highly conserved bipartite nuclear localization signal (NLS). This translocation is facilitated by the classical importin pathway. Notably, we demonstrate through reverse genetics that mutating this NLS prevents nuclear entry, and eliminates viral viability, confirming that the NLS is an active signal and that NS1 nuclear translocation is crucial for productive infection. In the nucleus, NS1 interacts with nuclear lamin A/C and chromatin, functioning as a transcriptional regulator by suppressing host genes involved in nucleic acid processing and pro-inflammatory responses, while activating a specific subset of antiviral genes. These findings uncover a previously unrecognized nuclear phase of DENV NS1 necessary for the full replication cycle of the virus.

## Materials and Methods

### Cell cultures, virus strain, virus propagation, and virus titration

Baby hamster kidney cells (BHK-21, ATCC CCL-10), Vero E6 (ATCC CRL-1586), and C6/36 *Aedes albopictus* cells (ATCC CRL-1660) were grown on MEM (Gibco, 41500-034) supplemented with 5% fetal bovine serum (FBS, Biowest, S1650) and 100 U/mL penicillin–streptomycin. Cultures were maintained at 37 °C for BHK-21 and Vero E6, and 28 °C for C6/36 cells in a water-saturated atmosphere with 5% CO2. DENV2, strain New Guinea, was harvested from infected suckling CD-1 mouse brains, passaged once in confluent C6/36 cells, and titrated in BHK-21cells by a plaque-forming assay. Briefly, tenfold serial dilutions from viral stocks or experimental supernatants were added to cell monolayers grown in 24-well plates. Virus absorption was allowed for 2h at 37°C in serum-free medium; afterward, the inoculum was removed, and the overlay medium was added (1X MEM, 5% FBS, and 0.75% carboxymethyl cellulose final). Five days after infection, the overlay medium was removed, cells were fixed with 10% formaldehyde for 10 minutes, and stained with 0.1% crystal violet solution for 30 minutes. After washing with tap water, plaques were counted and results expressed as PFU/ml. Procedures for suckling mice use were approved by our institutional ad hoc committee and performed following the Mexican official regulatory guideline NOM-062-ZOO-1999.

### Kinetics of NS1 nuclear localization in infected cells

Confluent BHK-21 monolayers, grown on glass coverslips in 24-well plates, were infected with DENV2 at a MOI of 3. At 6, 12, or 24 hpi, cells were fixed with 4% paraformaldehyde and permeabilized with 0.2% Triton X-100, with each step carried out for 10 min at room temperature. For NS1 detection by immunostaining, an anti-panflavi-NS1 mouse Mab (Abcam, AB214337) was used as a primary antibody, followed by an anti-mouse Alexa 488 conjugate (Invitrogen A-11001). Nuclei were counterstained with DAPI (Sigma Aldrich, D9542), and coverslips were mounted with Vectashield (Vector Laboratories, H-1000-10). The slides were analyzed using a Carl Zeiss LSM 900 confocal microscope, and the images were processed using ImageJ.

### Image deconvolution analysis

Two different deconvolution methods were used to quantify the DENV NS1 protein present in the nucleus of infected cells. In the first method, Z-stacks in which NS1 (Alexa488) and DAPI were detected were acquired. Regions of interest (ROIs) were manually segmented in the selected sections. The total cell area was defined by the plasma membrane, and the nuclear area by the DAPI staining boundaries. Quantification was performed using the spot detection tool of *ICY Bioimage Software,* with a detection scale of 1 to 3 pixels. The percentage of nuclear NS1 was calculated per cell as follows: % nuclear NS1 = (nuclear spots/total spots) x 100. The second method, based on *Volume Image Deconvolution Analysis (Fiji/ImageJ),* was developed in-house and used to strengthen the results. First, individual channels were extracted from the raw Z-stacks to generate a binary three-dimensional mask using the colocalization imaging plugin. The NS1 channel (Alexa488) was processed using a 3D median filter (filter = 1 pixel) followed by a high-pass filter (filter = 3 pixels). Nuclear edges were defined using the colocalization imaging plugin to generate a three-dimensional mask based on the DAPI signal. To isolate the NS1 nuclear signal, the masks and the NS1 channel (Alexa488) were processed, creating a dataset containing only the NS1 nuclear signal. A 3D object counter was used to quantify the number and density of NS1 clusters and perform a volumetric comparison between the total NS1 and the nuclear NS1. Statistical analyses and data visualization were performed in all cases using GraphPad Prism 8 for Windows. The results were expressed as the percentage of total intracellular NS1 present in the nucleus at a given time.

### In situ subcellular fractionation

In situ subcellular fractionation was performed using DENV2-infected cells at 24 hpi, following the protocol described by Sawasdichai et al. (27) with some modifications. Cells were incubated with cytoskeleton buffer (CSK; 10 mM PIPES pH 6.8; 300 mM sucrose; 100 mM NaCl; 3 mM MgCl_2_; 1 mM EGTA) supplemented with different concentrations of Triton X-100 to reveal distinct subcellular compartments. For **tightly held nuclear fractions**, whole cells were incubated with CSK buffer containing 0.1% (v/v) Triton X-100 for 2 minutes on ice to remove soluble cytoplasmic proteins. For the **chromatin fraction**, the remaining cellular structures were treated with CSK buffer containing 0.5% (v/v) Triton X-100 for 25 minutes on ice. To isolate the **nuclear matrix**, chromatin obtained from the previous step was digested by incubating the samples with CSK buffer supplemented with 100μg/mL of DNase I for 40 minutes at 37°C. After each extraction step, coverslips were washed with PBS and fixed with 4% formaldehyde for 12 minutes at room temperature. The efficiency of the fractionation procedure was validated by monitoring specific subcellular markers using primary antibodies against β-tubulin (Sigma-Aldrich, T7816), histone H2B (chromatin; ABclonal, A19812), and lamin B1 (nuclear lamina/matrix; Abcam, ab16048), followed by detection with Alexa Fluor 488 (Invitrogen, A-11001) and Alexa Fluor 647 (Invitrogen, A21245) conjugated secondary antibodies. NS1 was detected using the pan-flavivirus anti-NS1 Mab and secondary antibodies described above. Nuclei were counterstained with DAPI.

### Differential subcellular fractionation

Biochemical cell fractionation was performed using the subcellular protein fractionation kit (Thermo Scientific, 78840) following the manufacturer’s instructions. Confluent monolayers of BHK-21 cells grown in T25 flasks (approximately 6 x 10^6^ cells) were infected with DENV2 (MOI = 3), harvested at 24hpi, and treated with different cell lysis buffers to isolate cytoplasmic, membrane, nuclear soluble, chromatin-bound, and cytoskeletal protein fractions. All extraction buffers were supplemented with a protease inhibitor before use, and chromatin-bound nuclear proteins were recovered after incubation with micrococcal nuclease and CaCl_2_ (both provided with the kit) for 15 min at room temperature. The presence of NS1 in each fraction was determined by Western blot using an in-house-generated, rabbit anti-NS1 hyperimmune serum (7), and an anti-rabbit conjugated with horse radish peroxidase (Biorad, 172-1019), followed by detection with SuperSignal West Femto Maximum Sensitivity Substrate (Thermo Scientific, 34096) and visualization using a Newton 7.0 imaging system (Vilber). Quality and purity controls for each fraction were also determined by Western blot using the following antibodies; anti-HSP90 (Abcam, ab2927) for the cytoplasmic fraction, anti-calreticulin (Abcam, EPR3924) for the membrane fraction, anti-TFE3 (Invitrogen, PA5-54409) for the soluble nuclear fraction, anti-histone H3 (Santa Cruz, SC-517576) for the chromatin-bound fraction, and anti-vimentin antibody (Sigma-Aldrich, SAB4300676) for the cytoskeleton-bound fraction. All antibodies were used at the concentration range recommended by the manufacturers.

### Blue native polyacrylamide gel electrophoresis

To analyze NS1 protein complexes in their native state in the samples recovered from the differential subcellular fractionation, non-denaturing blue native PAGE (28), followed by Western blots, was carried out. Briefly, samples were prepared by mixing the protein fraction with 2X sample solution (0.187 M Tris-HCl, pH 6.8; 30% glycerol, and 80 μg/mL bromophenol blue) and Coomassie brilliant blue G-250 solution. The Coomassie dye was used to introduce a negative charge to the protein complexes without disrupting them, thereby facilitating separation based on the molecular mass of the complexes. The presence of NS1 was assessed by Western blot.

### Colocalization assays in infected cells

Confluent BHK-21 cell monolayers, grown on glass coverslips in 24-well plates, were infected with DENV2 at an MOI of 3, fixed at 24 hpi and processed for confocal microscopy. Cells were washed with PBS-0.2% Tween 20 and permeabilized with 300 μL of PBS-0.3% Triton X-100 for 10 minutes at room temperature. Afterward, the coverslips were transferred to a humid chamber over a sheet of Parafilm®, and 50 μl of blocking solution (glycine 25 mg/mL, 1% FBS, and 1% fish gelatin) was added for 2 h at 37 °C. The blocking solution was removed, and solutions containing an anti-panflavi NS1 mouse Mab, and an anti-lamin B1 Pab (Abcam, ab16048) as primary antibodies were added overnight at 4 °C. The solution with antibodies was removed, and the slides were washed 3 times for 10 minutes with PBS 0.2%-Tween 20, then 5 μL of a solution with secondary antibodies was added for 2 hours at 37 °C. Finally, the solution with secondary antibodies was removed, and a solution with DAPI (Merck, D9542) at 1:1000 was added for 10 minutes, and the slides were washed 3 times for 10 minutes with 0.2% PBS-Tween 20 and once for 5 minutes with Milli-Q water. The slides were mounted with 5μl of VectaShield (Vector, H-1000) and analyzed on an LSM 900 confocal microscope. For protein colocalization, approximately 100 cells were counted, and the Pearsońs and Manders correlation coefficients were calculated using the ICY software.

### Immunoprecipitation assays from transfected cells

Immunoprecipitation assays using NS1 as bait were carried out in BHK-21 cells transfected with 1 μg of a plasmid expressing DENV2 NS1 (a kind gift from Sebastián Aguirre and Ana Fernández Sesma, Icahn School of Medicine, Mount Sinai, USA), using Lipofectamine 3000 (Thermo Scientific, L3000015) according to the manufactureŕs instructions. The expressed protein also includes 28 amino acids corresponding to the E protein stem region for better stability and functionality, and an HA tag. At 24 hours post-transfection (hpt), cells were washed with ice-cold PBS and lysed using IP lysis buffer (50 mM Tris-HCl; 150 mM NaCl; 1% Triton X-100 supplemented with protease inhibitors (Merck, 11836170001). Immunoprecipitation was performed using the Pierce HA-Tag IP/Co-IP kit (Thermo Scientific, 26180) according to the manufactureŕs instructions. Proteins were eluted by resuspending the samples in 25 μL of 2X non-reducing sample buffer and heating at 95 °C for 5 minutes. Eluted proteins were separated by SDS-PAGE and analyzed by Western blots for the detection of NS1 and lamin A/C, and laminin B1. Primary antibodies were an anti-panflavi NS1, in-house-made, rabbit anti-NS1 hyperimmune serum, an anti-lamin A/C rabbit polyclonal antibody (ABclonal, A17319), and an anti-lamin B1 polyclonal antibody (Abcam, ab16048).

### Exogenous NS1 internalization assays

BHK-21 cells were incubated with commercial recombinant DENV2 NS1 protein (R&D Systems, 9439-DG) diluted in EMEM supplemented with 5% fetal bovine serum to a final concentration of 2 μg/mL. Cells were incubated for 6, 12, and 24 hours at 37°C. Following incubation, cells were fixed, and the intracellular localization of NS1 was analyzed by confocal fluorescence microscopy as described previously.

### Intracellular retention of NS1 assays

BHK-21 cells transfected with 1 μg of the plasmid encoding NS1 were treated at 6 hpt with brefeldin-A (BFA; Sigma-Aldrich; B7651), an inhibitor of the classical constitutive secretion pathway, at a final concentration of 3 μg/ml. Cells were fixed at 6 or 12 hours post-treatment and processed to evaluate the presence of NS1 inside the nucleus by confocal microscopy. To assess BFA treatment efficiency, the presence or absence of NS1 in cell supernatants was determined using lateral flow immunoassays specific for DENV NS1 (a kind gift from Irene Bosch, Massachusetts Institute of Technology, USA).

### Nuclear Localization Signal (NLS) identification

Potential (NLS) in the DENV2 NS1 sequence from strain New Guinea C (GenBank: AAC59275.1) were analyzed using the NLS Mapper Predictor software (29). The analysis identified a putative bipartite NLS at residue 189 with a score of 5.9 (where 1.0 indicates total cytoplasmic localization, and 10.0 total nuclear localization), thus predicting an NS1 localization in both the nucleus and cytoplasm.

### Sequence alignment and conservation analysis

NS1 amino acid sequences from the four DENV serotypes and Zika virus were extracted from the NCBI GenBank database (accession numbers: NP_059433.1 for DENV1, AAC59275.1 for DENV2, YP_001621843.1 for DENV3, NP_073286.1 for DENV4, and YP_002790881.1 for ZIKV). Multiple sequence alignment was performed and visualized using the Jalview software (30) to assess the degree of conservation, the putative bipartite NLS region (residues 189-216), and the adjacent N-linked glycosylation site (N207) across the analyzed orthoflavivirus.

### Structural modeling of NS1 and NLS location

Three-dimensional structures of the dimer and hexamer of DENV2 NS1 (GenBank: AAC59275.1) were predicted using AlphaFold 3.0 (31). The resulting models were subsequently used to generate molecular graphics to map the NLS location in the 3D structure of NS1 (32) and visualize its spatial distribution.

### Recombinant pNS1 and NLS mutations

To evaluate the functionality of the predicted NLS, the sequence was mutated by site-directed mutagenesis on the NS1-expressing plasmid. The NS1 coding sequence was excised using SacI (NEB, R0156) and PvuII (NEB, R0151S) restriction enzymes, and the fragment was used as a template for a modified overlap extension PCR (33). Basic residues within the predicted NLS were substituted with glutamine (Q). Three mutants were constructed: Mut1, containing substitutions K_188_Q and R_191_Q; Mut2, containing substitutions K_210_Q and K_213_Q, and Mut3, combining both mutations. The mutated fragments were ligated back into the vector using T4-ligase (NEB, M0202S). Finally, the integrity of all constructions was verified using Sanger sequencing (Macrogen, Inc., Seoul, Korea). Primers used for mutagenesis and sequencing are listed in Supplemental Table 1.

### Reverse genetics and rescue of recombinant viruses

The NLS mutations K_188_Q, R_191_Q, K_210_Q, and K_213_Q were all introduced into a DENV2 infectious cDNA (icDNA) clone harboring mCherry marker (34). Recombinant icDNA plasmids (DENV2-wt-mCherry and DENV2-mut-mCherry) were linearized using the AgeI restriction enzyme (New England Biolabs, R0552S). After linearization, the templates were *in vitro* transcribed (IVT) with the mMessage mMachine SP6 transcription kit (Invitrogen, AM1340) to generate 5’-capped viral RNAs, following the manufactureŕs recommendations. To perform rescue, Vero E6 cells were transfected with the IVT RNAs using Lipofectamine LTX reagent (Thermo Fisher Scientific, 15338030). Transfected cells were maintained in 5 mL of culture medium supplemented with 5% FBS. Viral replication was monitored daily by observing the expression of the mCherry fluorescent reporter. After 8 days of transfection, the culture supernatant (P0 stocks) was collected and clarified by centrifugation. The infectious titers of P0 were quantified by plaque assay in BHK-21 cells. Viral stocks derived from the icDNA clones were maintained by directly inoculating BHK-21 monolayers with P0 supernatants.

### Rescue of mutant virus using trans-complementation

Trans complementation assay of DENV2-mut-mCherry was performed as follows. BHK-21 cells were grown to approximately 60% confluence in 24 wells plates, and transfected using Lipofectamine LTX, according to the following conditions: i) non-transfected cells (negative control); ii) cells transfected with 1 μL of IVT DENV2-wt-mCherry RNA (wt-RNA control); iii) cells transfected with 1 μl of IVT DENV2-mut-mCherry RNA (mut-RNA control); iv) cells transfected with 500 ng of wt-NS1-expressing plasmid (plasmid control); v) cells co-transfected with 1 μL of IVT DENV2-mut-mCherry RNA + 500 ng of wt-NS1 expressing plasmid (pNS1-wt); and vi) cells co-transfected with 1 μL of IVT DENV2-mut-mCherry RNA + 500 ng of mutant Mut3-NS1 expressing plasmid.

To ensure experimental uniformity, transfection complexes were individually prepared for each genetic material (IVT RNA and plasmid) before being mixed in the well. This strategy ensured that the total amount of Lipofectamine LTX reagent, nucleic acids, and volume remained constant relative to single-type nucleic acid transfection controls. Rescue efficiency was evaluated 6 days post-transfection by visual assessment of mCherry intracellular expression and by evaluating virus progeny production.

### Transcriptomic profiling of BHK-21 cells expressing wild-type and NLS-mutant DENV NS1

BHK-21 cells were cultured in 6-well plates and grown to approximately 80% confluence. Cells were transfected with 1.5 μg of plasmid DNA per well using Lipofectamine 3000 transfection reagent (Thermo Scientific), following the manufactureŕs instructions. Three experimental conditions were used: i) empty vector (negative control); ii) plasmid expressing wild-type NS1 (wt-NS1); and iii) plasmid expressing NS1 with the NLS mutated (Mut3-NS1).

Transfections were performed in independent duplicate wells for each condition to generate biological replicates. Successful expression of NS1 was qualitatively validated using lateral flow immunoassays 24 hpt. To minimize well-to-well variability and ensure biological representativeness, the two independent biological replicates for each condition were pooled before RNA extraction. Total RNA was extracted from the samples using the RNeasy Mini Kit (Qiagen, 74104) following the manufactureŕs instructions and incorporating a DNA digestion step to remove residual plasmid DNA. The concentration and purity of the RNA were measured using a Nano Drop spectrophotometer (Thermo Fisher). The RNA-seq library construction, high-throughput sequencing, and preliminary bioinformatic analysis were carried out by a commercial service provider (Macrogen, Seoul, Korea). In brief, RNA integrity was evaluated using the Agilent 4200 TapeStation system; the sequencing libraries were generated using the TruSeq Stranded Total RNA with Ribo-Zero Gold Kit (Illumina) to efficiently deplete cytoplasmic and mitochondrial ribosomal RNA while strictly preserving transcript strandedness. Library integrity and fragment size distribution, exhibiting a characteristic peak spanning 320–338 bp, were rigorously assessed using the Agilent 2100 Bioanalyzer. Subsequent library quantification was performed via quantitative PCR (qPCR) to ensure optimal cluster generation. The validated libraries were subjected to high-throughput sequencing on the Illumina NovaSeq X platform. All raw sequence data in FASTQ format have been deposited in the NCBI Sequence Read Archive (SRA) under the accession number PRJNA1445342.

The raw sequence data underwent an initial quality assessment using FastQC (v0.11.7) to evaluate per-base quality scores, adapter contamination, and GC content distribution. To ensure high-fidelity downstream analysis, low-quality bases and residual adapter sequences were systematically removed using Trimmomatic (v0.38). The resulting high-quality clean reads were aligned to the *Mesocricetus auratus* (golden hamster) reference genome (BCM_Maur_2.0) using the splice-aware aligner HISAT2 (v2.1.0) (35). Following alignment, transcript assembly and quantification of gene expression levels were performed using StringTie (v2.1.3b), which facilitated the estimation of transcript abundance for subsequent differential expression analysis.

### Bioinformatic analysis

Differential expression analysis was performed using the edgeR package, employing a generalized linear model (GLM) framework based on the negative binomial distribution. To account for compositional biases between libraries, raw read counts were normalized using the Trimmed Mean of M-values (TMM) method. Differentially expressed genes (DEGs) were strictly defined based on a biological effect size of absolute log2 fold change > 1 and a statistically significant threshold of p-value < 0.05.

To elucidate the biological processes perturbed by the NS1 mutation, Gene Ontology (GO) overrepresentation analysis was executed utilizing the **clusterProfiler** (36) R package. *Mesocricetus auratus* GO annotations were systematically retrieved via the **biomaRt** interface (37), querying the Ensembl database, and enrichment was conducted using the enricher () function. To quantify the magnitude of enrichment intensity for each GO term, a standardized Z-score was computed. This metric was calculated by evaluating the deviation of the observed number of term-associated DEGs from the expected frequency, normalized by the square root of the expected value. The expected frequency was derived from the background proportion of term-annotated genes relative to the total genomic universe, scaled by the input DEG subset size. Enriched GO terms were systematically stratified into five overarching functional modules based on semantic similarity and keyword association: Immune Response, Inflammation, Lipid Metabolism, Energy/Mitochondrial Function, and Nucleic Acid Metabolism.

### Statistical analysis

Results were analyzed for statistical difference by 2-way ANOVA using GraphPad Prism version 9.0.

## Results

### DENV2 NS1 translocates to the nucleus and associates with chromatin and nuclear lamina

DENV NS1 is usually described as an ER-resident or secreted glycoprotein. Here based on preceding indirect evidence, we investigated its potential subcellular distribution in other compartments. Using confocal immunofluorescence microscopy, an accumulation of NS1 in the nucleus was observed in infected BHK-21 cells (Figure 1A). Time-course analysis revealed that nuclear translocation is time-dependent, with nuclear signal detectable as early as 6 hours post-infection (hpi), and increasing up to 24 hpi. (Figure 1A). Quantitative analysis using both ICY and ImageJ software indicated that up to 30% of the total intracellular NS1 pool can be present in the nucleus at 24 hpi (Figure 1B).

**Figure 1.**
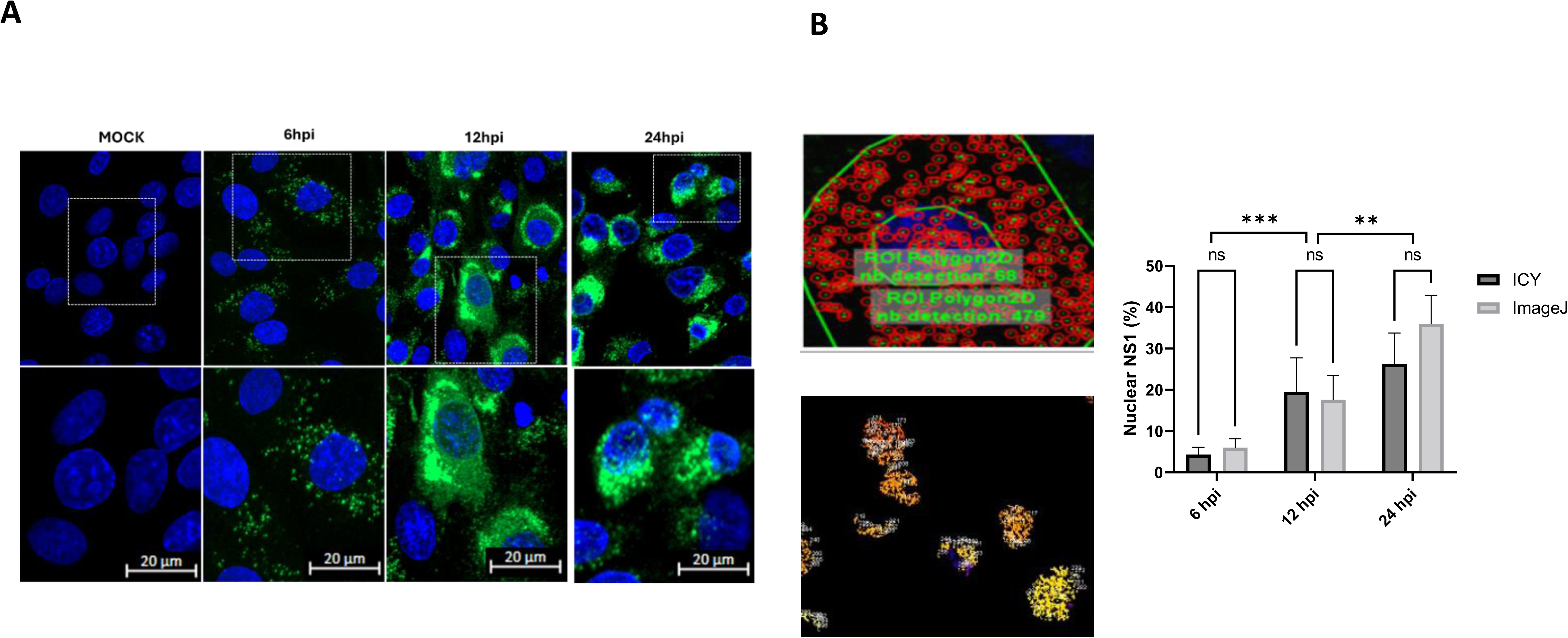
Nuclear localization of DENV-2 NS1 protein. **A)** BHK-21 cells were infected with DENV-2, fixed at the indicated times post-infection, and processed for confocal microscopy. NS1 (green) and nuclei (DAPI, blue). **B)** Quantification by two different bioinformatics software, ICY © (upper panel and ImageJ (lower panel), of the amount of NS1 present inside the cell nuclei. Results are expressed as a percentage, taking as 100% the total pool of NS1 present inside the cell (right panels). **/*** *p=0.005.* There are no significant differences between the two methods

To better define the sub-nuclear localization of NS1, we performed “*in situ*” subcellular fractionation. Images showed that NS1 was found not only in soluble form within the nucleoplasm but also retained in the chromatin-bound fraction and the nuclear matrix (Figure 2A). Quality controls validated the fractionation process and, therefore, the nuclear signal observed for NS1 (Figure 2B). Cell biochemical fractionation corroborated the in situ fractionation findings, detecting NS1 in the soluble nuclear, chromatin-bound, and cytoskeletal fractions of infected cells (Figure 3A). Furthermore, blue native PAGE (BN-PAGE) analysis of these fractions indicated that nuclear NS1 may exist as dimers and possibly as tetrameric complexes (Figure 3B). Quality control of the fractions using different protein markers suggested that the fractions were clean (Figure 3C), supporting the results indicating the presence of different forms of NS1 inside the nucleus.

**Figure 2.**
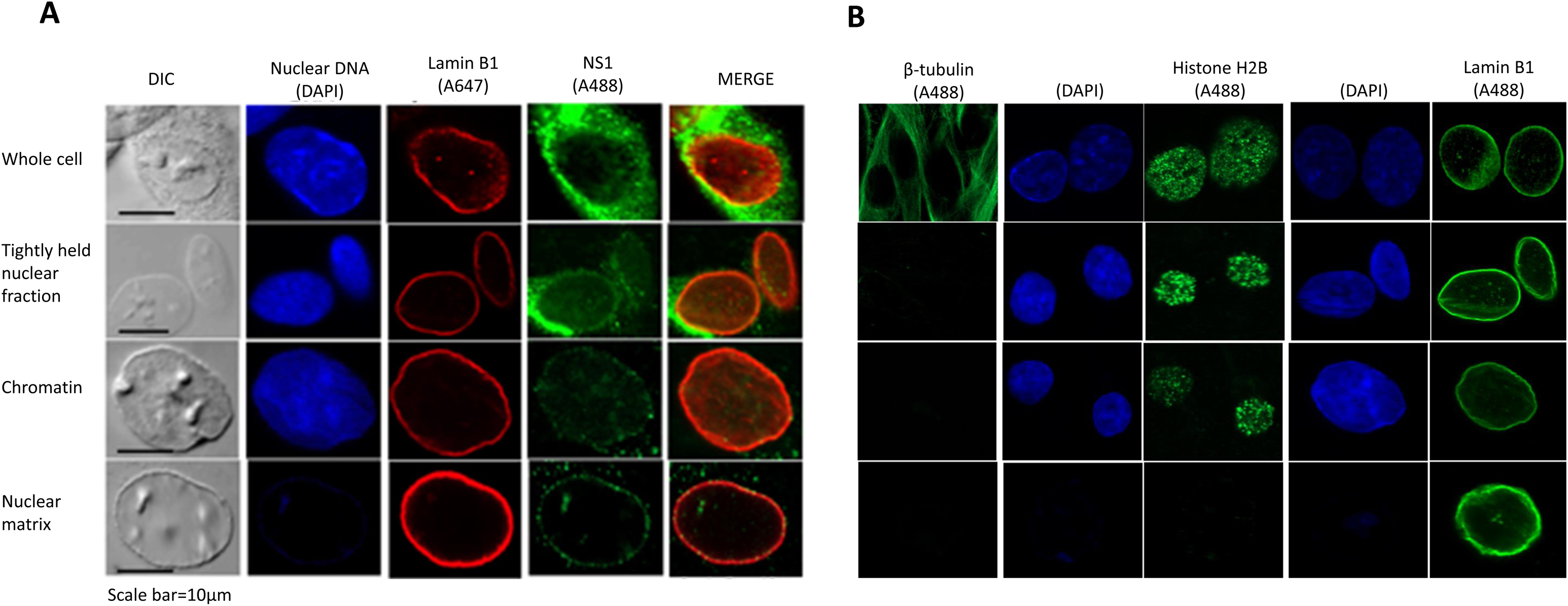
Analysis of the subcellular localization of NS1 by in situ cell fractionation. BHK-21 cells were infected with DENV-2, fixed at 24 hpi, and examined using confocal microscopy. **A) C**olocalization of NS1 with cell structures of interest. **B)** Labelling of different cell proteins for fraction quality control. Sequential removal of different cell parts was achieved throughout the process based on buffer compositions. DIC, differential interference contrast. Results from one of two independent experiments are shown.

**Figure 3.**
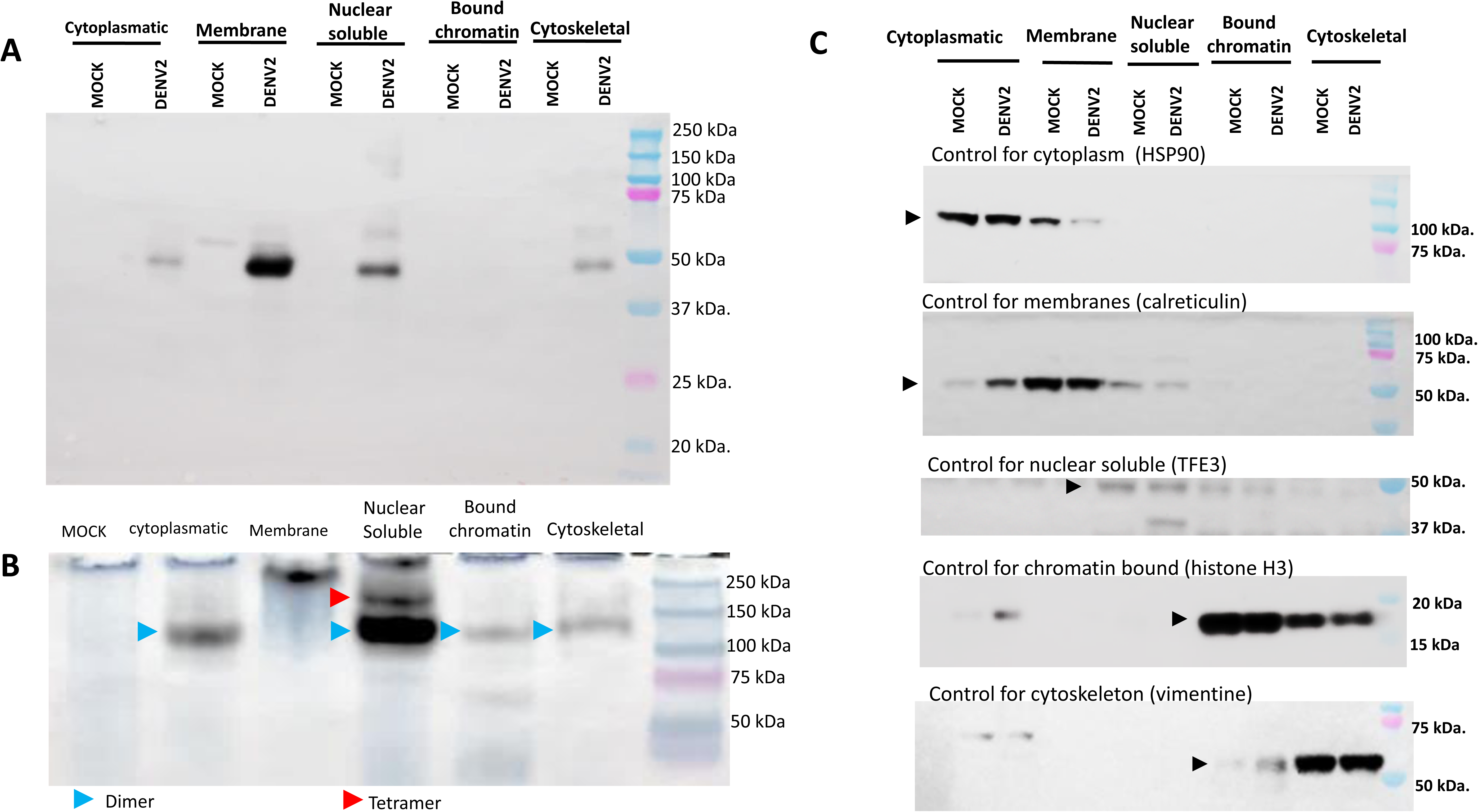
Analysis of the subcellular localization of NS1 by cell fractionation. BHK-21 cells were infected with DENV2 or mock**-**infected, harvested at 24 hpi, and subjected to subcellular fractionation. **(A)** Localization of DENV2 NS1 by western blot after fully denaturing SDS-PAGE electrophoresis, or **(B)** after non-denaturing blue native PAGE electrophoresis. **(C)** Western blot for quality control of each fraction using appropriate protein markers. Results from one of two independent experiments are shown.

The fluorescence patterns obtained for NS1 after the in-situ fractionation (Figure 2A), especially in the nuclear matrix fraction, suggest a location for NS1 similar to that of lamin, around the nuclear envelope. Moreover, a certain degree of colocalization between NS1 and lamin was observed. In order to verify this interaction, enhanced confocal analysis and co-immunoprecipitation (co-IP) assays were carried out using infected cells and transfected cells, respectively. Confocal images suggest strong colocalization between NS1 and lamin B1 at 24 hpi (Figure 4A). The physical interaction with the nuclear envelope was further validated by co-IP assays, which show that NS1 interacts directly with lamin A/C and lamin B1 (Figure 4B). To rule out artifacts due to variations in input protein abundance, densitometric IP/Input ratios were calculated. This normalization confirmed a specific enrichment of lamin proteins in the presence of NS1, with Lamin A/C showing a strong increase in co-immunoprecipitation (∼7–12-fold across replicates) and Lamin B1 a more moderate enrichment (∼3.3-fold) compared to non-transfected controls. Together, these data demonstrate that part of NS1 translocates to the nucleus during DENV infection, where it associates with key structural components.

**Figure 4.**
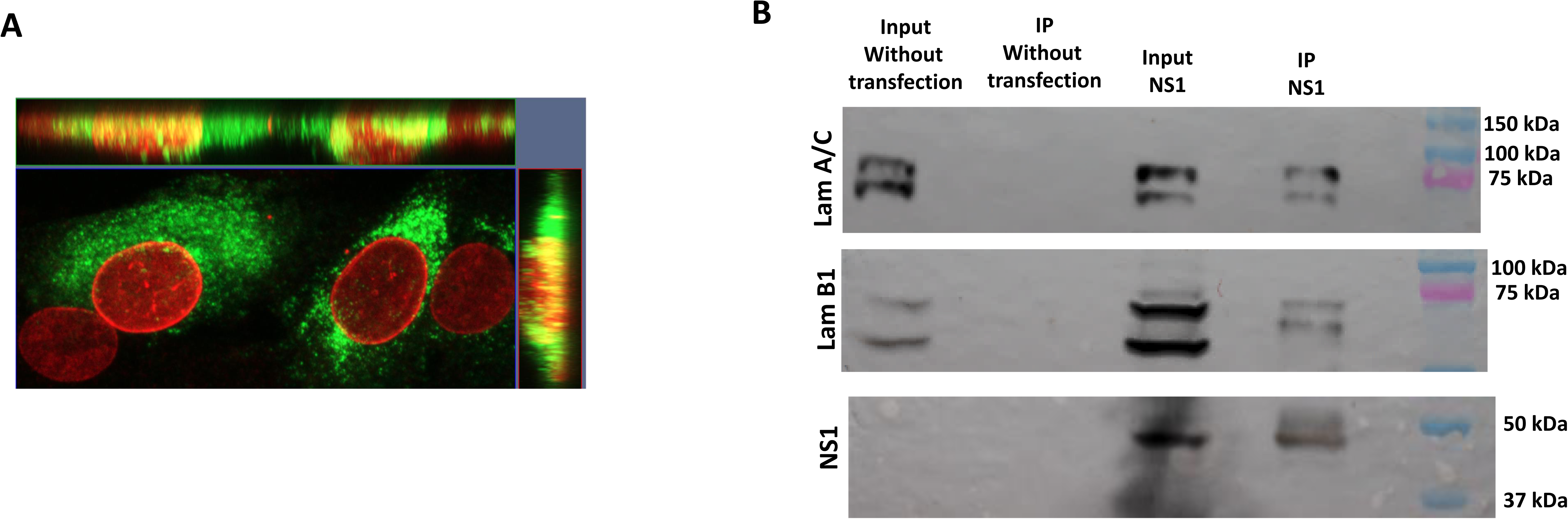
Interaction of the DENV2 NS1 with nuclear lamin. **(A)** BHK-21 cells were infected with DENV2 and processed at 24 hpi. Colocalization was analyzed by confocal microscopy, detailing the spatial interaction between NS1 (green) and the nuclear lamina (red). Signal overlap was quantified by Pearson’s (0.162) and Manders (0.324) correlation coefficients. **(B)** Co-immunoprecipitation assay from BHK-21 cells transfected with a plasmid expressing DENV2 NS1. Cells were collected at 24 hpt and precipitation was performed using NS1 as a bait. Results from one of two independent experiments are shown.

We also evaluated whether endogenous, secreted, or both populations of NS1 could reach the cell nucleus. Uninfected BHK-21 cells exposed to soluble NS1 and analyzed by confocal microscopy at 6, 12, and 24 h after exposure showed the presence of NS1 inside the nucleus at all time points after treatment (Supplemental Figure 1). In addition, DENV2-infected cells treated with BFA to prevent NS1 secretion also showed the presence of NS1 inside the nucleus at 6 and 12 h after treatment (Supplemental Figure 2). These results suggest that both intracellular and secreted NS1 can traffic to the cell nucleus.

### NS1 nuclear entry is mediated by a conserved bipartite NLS via the classical importin α/β importin pathway

The molecular mass of NS1 dimers (≈100 kDa) is too high to allow for the passive transport into the nucleus, as active transport is required for proteins larger than 40-50 kDa (38,39). Thus, to understand the mechanism of NS1 translocation, we analyzed the DENV2 NS1 sequence in search of NLS using cNLS Mapper software and identified a putative bipartite NLS located at residues 189-216 (sequence KDNRA-17X-KIEKAS), with a score of 5.9. The sequence is conserved across all four DENV serotypes and even in ZIKV (Figure 5A and B). Structural modeling using AlphaFold 3.0 suggests that the bipartite NLS is located in the β-strands 11, 12, and 13 of the ladder domain of NS1, flanking the glycosylated N207 position, and is exposed on the surface of the dimer and the hexamer; thus, it is theoretically accessible for interaction with nuclear transport receptors (Figure 5C).

**Figure 5.**
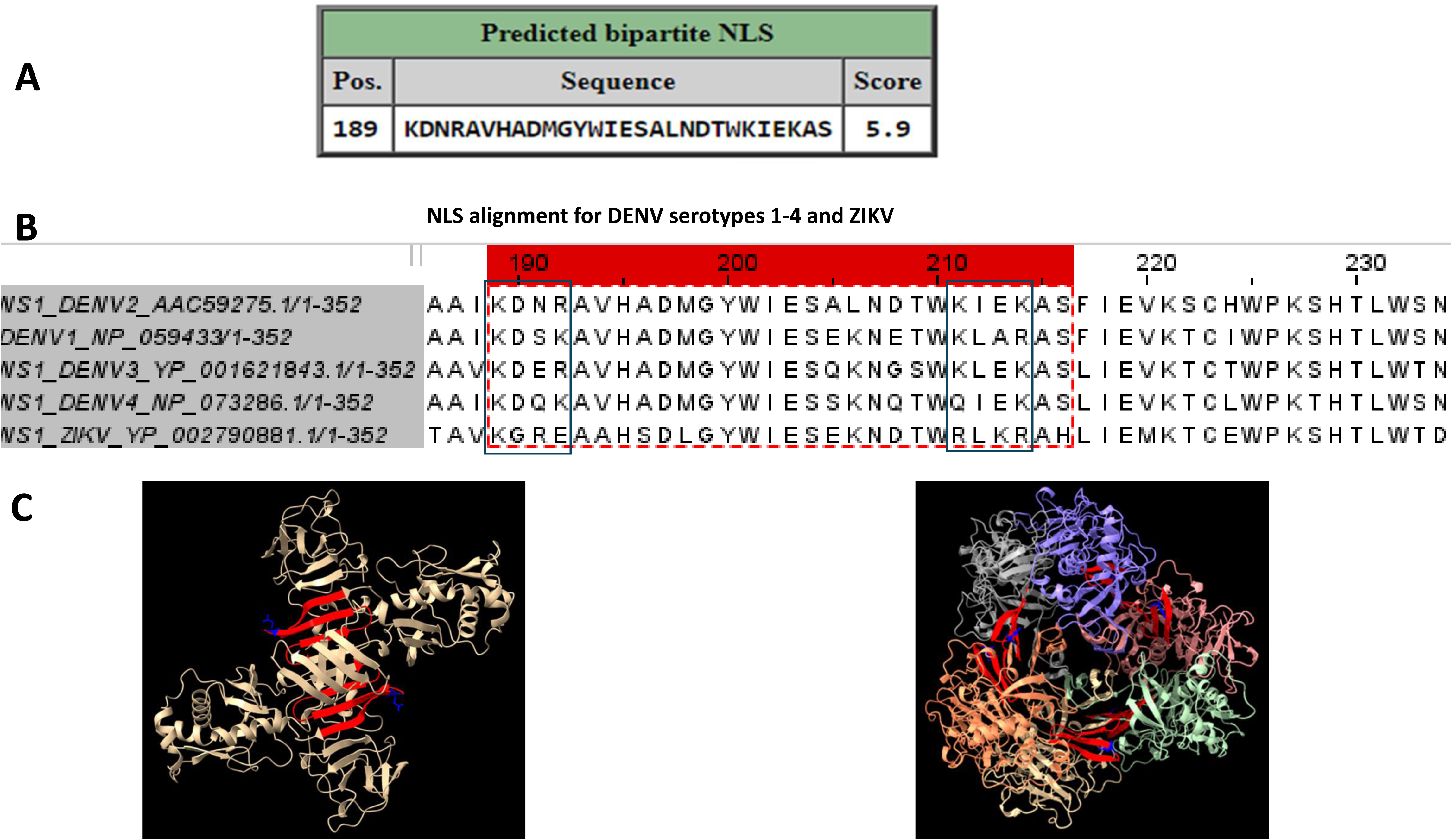
*In silico* identification and mapping of a nuclear localization signal (NLS) in DENV2 NS1 protein. **A)** Bipartite NLS sequence predicted in DENV-2 NS1 by NLS mapper (score of 5.9, with a total of 1.0 cytoplasmic location and 10.0 nuclear location). **B)** Sequence alignment of the predicted bipartite NLS across DENV serotypes 1 to 4 and Zika virus. **C)** Structural models of dimeric (left) and hexameric (right) DENV-2 NS1 generated by Alphafold 3.0 showing the exposed nature of the predicted NLS (highlighted in red and N-glycosylation site (N207) in blue).

Functional validation of the NLS was performed using site-directed mutagenesis. Mutants in the first (Mut1; K189Q/R192Q), second (Mut2; K211Q/K214Q), and both parts (Mut3) of the NLS were generated to neutralize the basic residues (Figure 6A). Western blot analyses were performed to assess the expression levels of the NS1 mutants relative to wt-NS1 and revealed no clear differences between the expression levels of wt-NS1 and Mut3-NS1 (Supplemental Figure 3). Immunofluorescence analysis of transfected BHK-21 cells showed that approximately 20% of the total wt-NS1 accumulated in the nucleus. No significant change was observed for Mut2 (Figure 6B and C). In contrast, a significant reduction in the amount of nuclear NS1 was observed for Mut1 and Mut3 (20% versus 15% of the total intracellular NS1 pool, *p<0.005*). Equal levels of reduction were observed in cells treated with 0.5µM ivermectin, a specific inhibitor of the importin α/β transporter. Control experiments carried out with the protein NLS-4-GFP suggested that the 0.5 µM ivermectin concentration used was effective (Figures 6B and 6C). These results establish that NS1 nuclear import is an active process driven by a functional bipartite NLS and dependent on the host classical nuclear import machinery.

**Figure 6.**
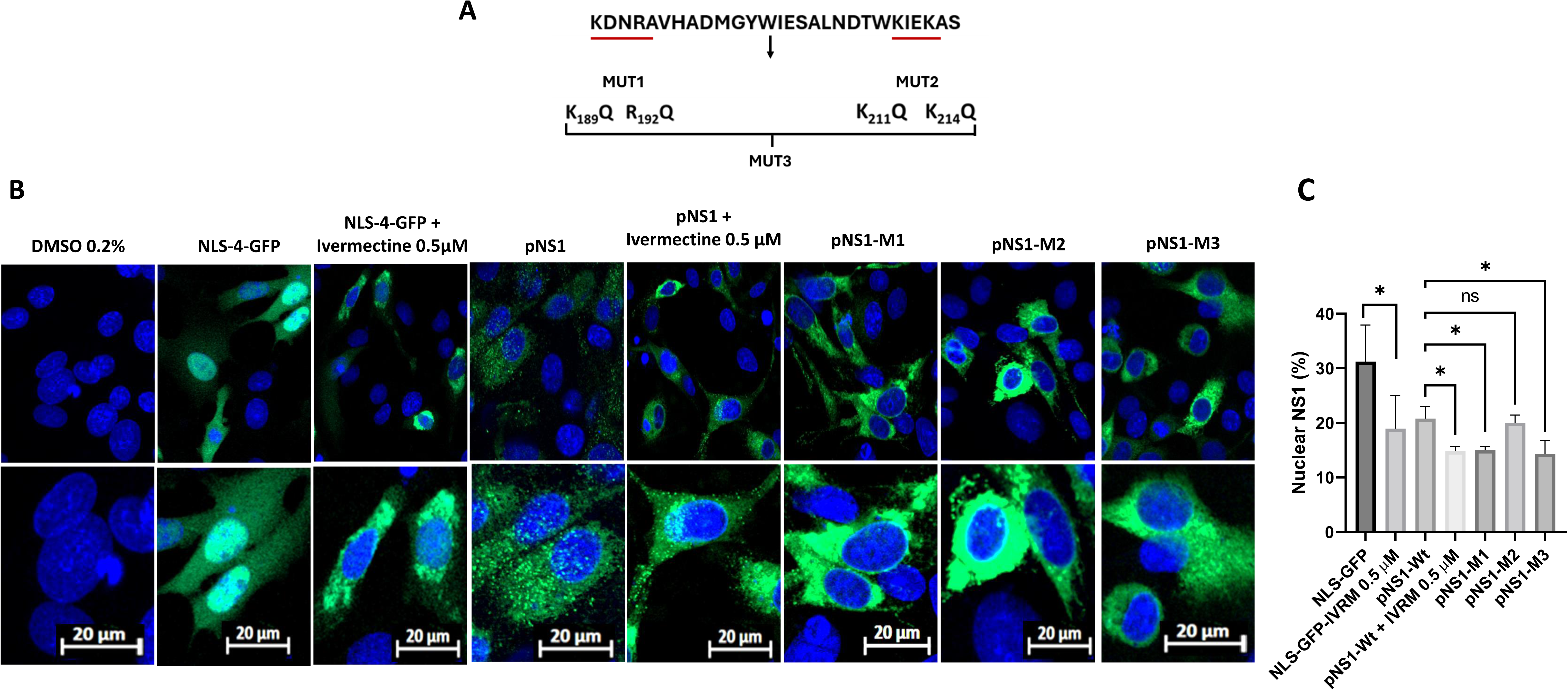
Experimental validation of the functionality of the bipartite NLS in the DENV-2 NS1 protein. **A)** Mutations introduced in part 1 (Mut1), part 2 (Mut2) or both parts (Mut3) of the bipartite NLS (underlined in red). Residue changes and positions are indicated. **B)** Evaluation of the amount of NS1 present in the cell nuclei after transfection with the different mutant constructions. Left panels, plasmid expressing GFP with 4 NLS (4-GFP NLS) used as a positive control. Right panels, wild type (pNS1) and the 3 different mutant constructions. Ivermectin treatment of 4-GFP NSL and pNS1-transfected cells was included as an additional positive control. Confluent monolayers of BHK-21 cells were processed for confocal microscopy 24 hpt. Cell nuclei were stained with DAPI (blue), and NS1 was visualized with Alexa488 (green). **C)** Quantification of the amount of NS1 present inside the cell nuclei, expressed as a percentage of the total signal. For each construction, 30 cells were analyzed. *p<0.05

**Figure 7.**
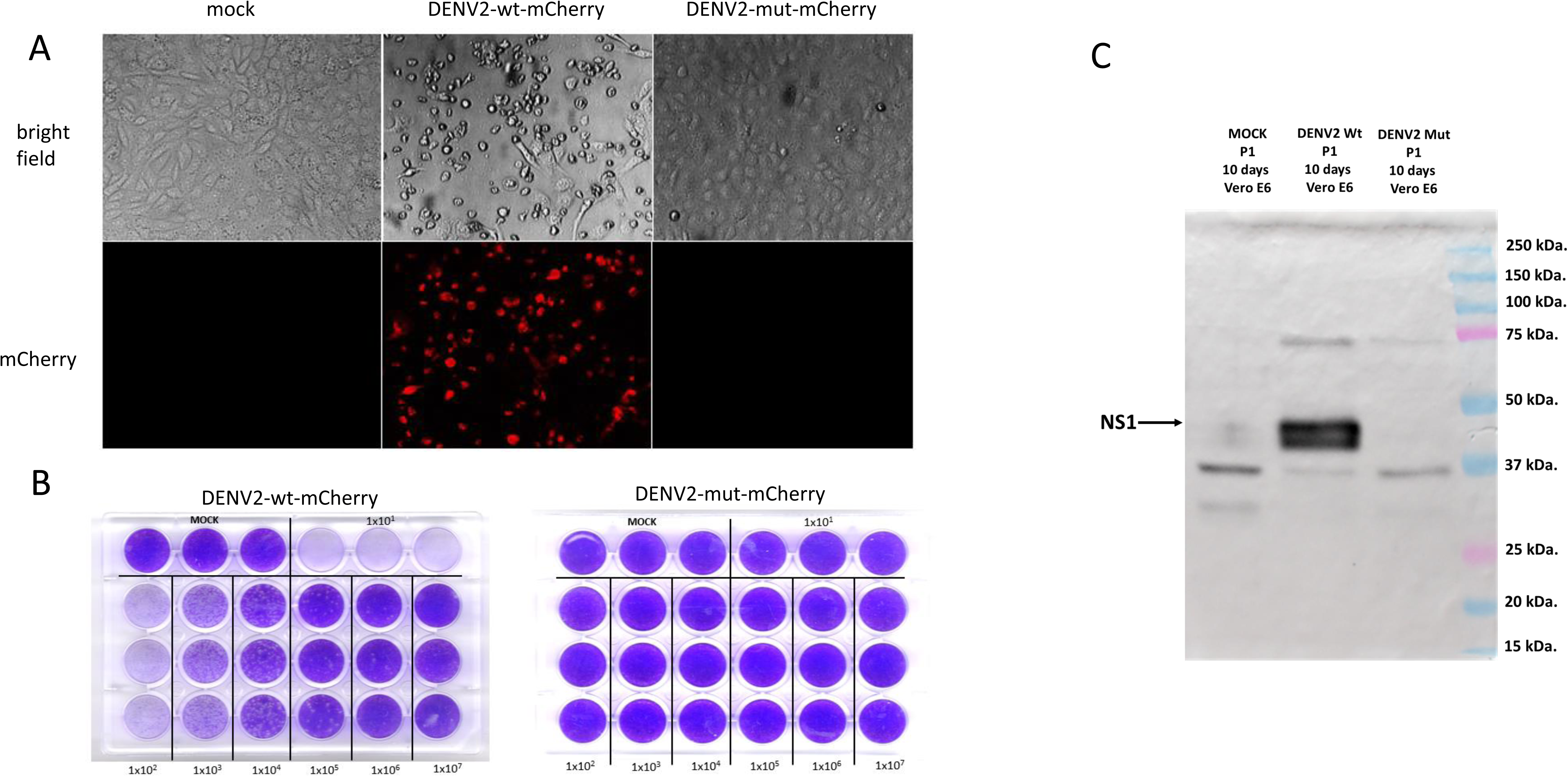
Characterization of DENV-2 wild-type and DENV-2 NLS mutant infectious clones. Vero E6 cells were transfected with in vitro transcription-derived RNA from infectious clones of DENV-2 wild type and DENV-2 NLS mutant, which also express mCherry as a marker, and cell supernatants were collected 6 days after transfection (passage 0). **A)** Epifluoresce assay of Vero E6 cells inoculated with passage 0 and fixed 6 days after inoculation. **B)** Titration in BHK-21 cells of cell supernatants collected from Vero E6 cells 6 days after inoculation (passage 1). Titer for Wt virus was calculated in 1.6×10^7^ PFU/ml. No plaque was observed for the mutant strain. **C)** Western blot analysis of NS1 protein expression in lysates from infected Vero E6 cells harvested 6 days after inoculation. Results from one of two independent experiments are shown.

### Nuclear translocation of NS1 is essential for DENV2 replication and infectious virion production

Experiments with BHK-21 cells infected with DENV and treated with ivermectin were inconclusive due to the generalized effect of ivermectin on DENV replication (Supplemental Figure 4) (40,41). To determine the biological relevance of nuclear NS1, an icDNA clone of DENV2, DENV2-mut-mCherry, carrying an mCherry reporter and harboring the same substitutions as Mut3-NS1, was constructed. In Vero E6 cells, transfection with transcripts of the DENV2-wt-mCherry revealed effective rescue and robust replication, as indicated by mCherry fluorescence and expression of NS1 detected by Western blot. In addition, infectious viral progeny was also detected (1×10^5^ PFU/mL). In contrast, no evidence of viral protein expression or virus progeny production was detected in the cells transfected with RNAs corresponding to the DENV2-mut-mCherry. These results indicate that the trafficking of NS1 to the nucleus during infection is necessary for a successful viral rescue and replication.

We hypothesized that the defect in the DENV2-mut-mCherry was due to the specific loss of nuclear NS1 functions rather than structural instability of the viral RNA or NS1 protein. To test this hypothesis and corroborate the need for NS1 to reach the cell nucleus to promote a successful viral replication, a trans-complementation assay in BHK-21 cells was carried out. Again, the transfection of cells with the DENV2-mut-mCherry RNA alone resulted in no replication. However, providing wt-NS1 in trans (via transfection with wt-NS1 plasmid) resulted in successful viral rescue, as evidenced by the recovery of mCherry expression (Figure 8A) and virus progeny production, as indicated by the formation of infectious foci in reinfection assays (Figure 8B). Interestingly, the trans-complementation with the Mut3-NS1 plasmid also resulted in virus rescue (Figure 8A); however, no infectious virus progeny was recovered in the supernatants (Figure 8B), an observation indicative of an abortive infection. These results indicate that the ability of the NS1 to enter the nucleus of infected cells is necessary for completion of a fully productive infection.

**Figure 8.**
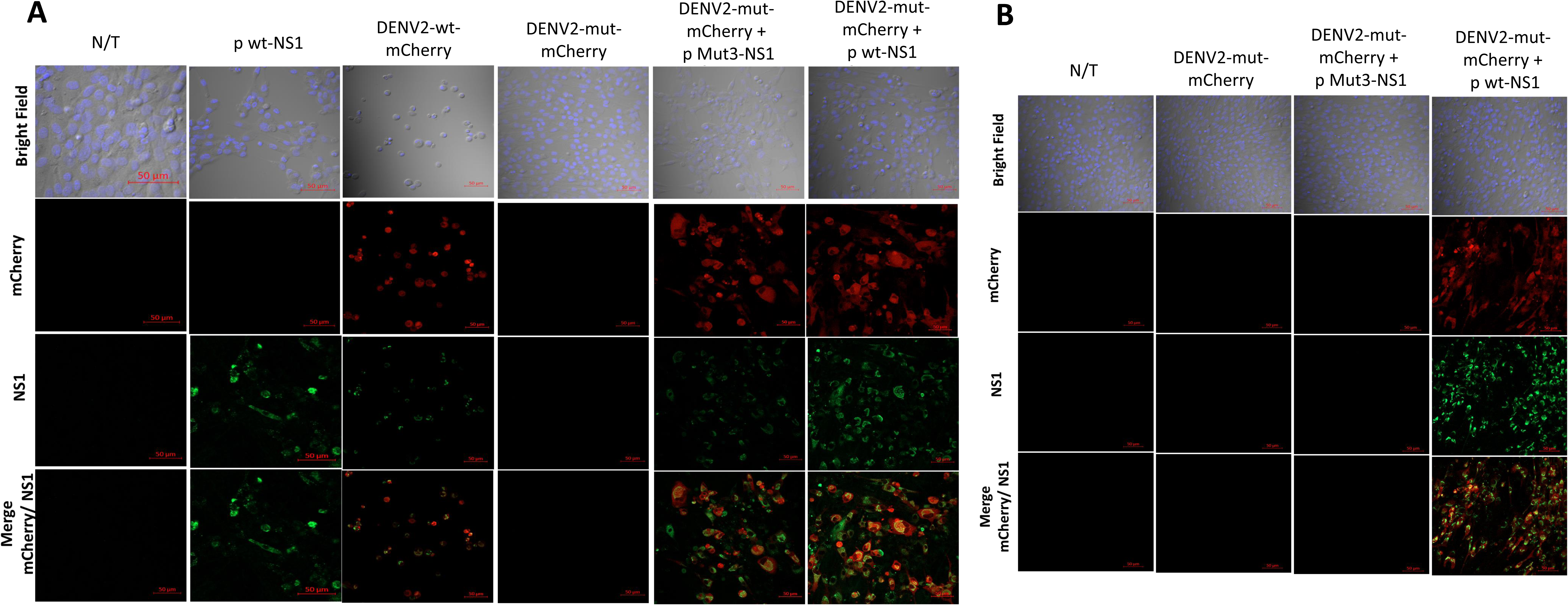
Rescue of DENV-2 NSL mutant infectivity by complementation in trans with NS1. **A)** Monolayers of BHK-21 were simultaneously transfected with in vitro transcribed RNA from DENV2-mut-mCherry, together with plasmids that express wild-type NS1 (p wt-NS1) or NS1 mutated in both parts of the NSL (p Mut3-NS1), and 6 days after, cells were fixed and processed for confocal microscopy. Controls include non-transfected cells (N/T), cells transfected with a plasmid expressing wt-NS1 wild type alone (p wt-NS1), and RNA derived from either infectious clone alone (DENV2-wt-mCherry and DENV2-mut-mCherry). **B)** Cell supernatants collected from the trans complemented cells after 6 days were tested for infectivity in monolayers of BHK-21 cells. Cells were fixed and processed for confocal microscopy 7 days after infection. Cell nuclei were stained with DAPI (blue), and DENV NS1 was visualized using an anti-NS1 Mab as primary antibody and an Alexa 488 conjugated secondary antibody (green). Scale bar: 50uM. Results from one of two independent experiments are shown.

### Nuclear NS1 modulates the cell transcriptional landscape

To further elucidate the functional implications of NS1 within the nucleus, we performed a comparative high-throughput RNA-seq transcriptomic profiling of BHK-21 cells expressing either wt-NS1 or the Mut3-NS1 variant. Cells transfected with an empty plasmid were included as a control. The robustness of the data set was confirmed by Pearsońs correlation analysis, where high correlation indexes (r > 0.98) between biological replicates were found in all three conditions (control, wt, and Mut3), indicating high reproducibility and data reliability (Supplemental Figure 5). Differential expression analysis revealed a deep shift in the transcriptome in BHK-21 cells depending on the subcellular localization of transiently expressed NS1.

When comparing cells expressing mutated NS1 against those expressing wt-NS1, a total of 1,596 differentially expressed genes (DEGs) were identified, with 1,052 genes downregulated and 544 upregulated significantly (Figure 9A and Supplemental Figure 6). This transcriptional variance is illustrated by the two-way hierarchical clustering heatmap, which shows a distinct expression profile that visibly separates the wt-NS1 effect from the Mut3-NS1 phenotype (Figure 9B). These results indicate significant changes in both the number and the nature of genes regulated when NS1 reaches the nucleus compared with when it is retained in the cytoplasm.

**Figure 9.**
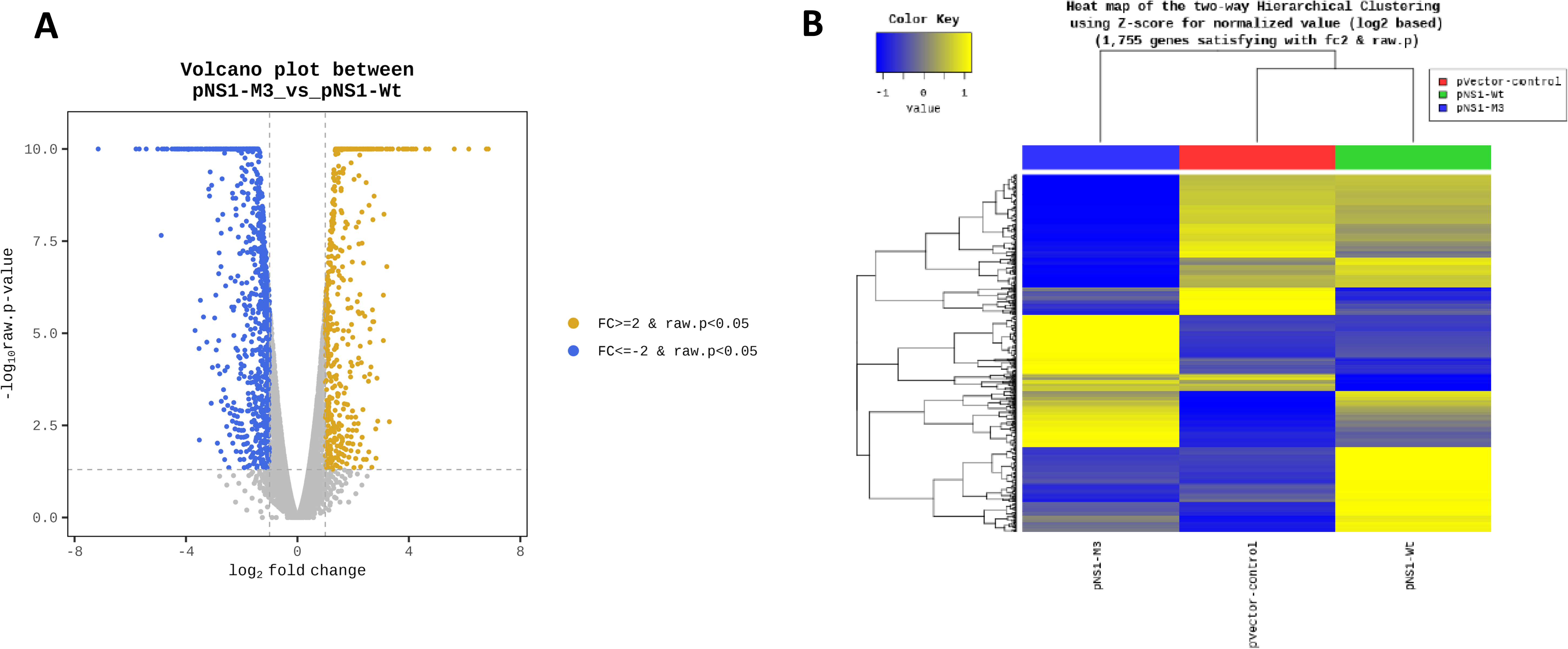
Direct transcriptomic comparison between cells expressing Mut3-NS1 (pNS1-M3) and wt-NS1 (pNS1-Wt) and hierarchical clustering analysis. **A)** Volcano plot showing differential gene expression in BHK-21 cells expressing the NS1-M3 mutant relative to wild-type NS1-Wt. The x-axis represents the fold change (log₂), and the y-axis indicates statistical significance (-log₁₀ crude P-value). Orange dots denote significantly upregulated genes in NS1-M3 compared to NS1-Wt, while blue dots represent downregulated genes (log₂ FC ≥ 1, P < 0.05). The extensive distribution of differentially expressed genes highlights the distinct biological impact of the M3 mutation compared to the native protein. **B)** Bidirectional hierarchical clustering heatmap of 1,755 genes with significant differential expression (times change > 2 and cutoff P-value). Rows represent individual genes and columns represent experimental conditions. Expression levels are presented as Z-scores of log2 normalized values, ranging from low expression (blue) to high expression (yellow). The upper dendrogram reveals a distinct clustering of the NS1-M3 mutant, separate from the NS1-Wt and control groups. (Right)

Gene Ontology (GO) enrichment analysis provided insights into the biological processes regulated by nuclear NS1. The 1,052 genes that were downregulated in the Mut3-NS1 condition but induced by wt-NS1 were strongly associated with nucleic acid metabolism (566 genes), suggesting that nuclear NS1 actively reprograms the host machinery to support viral RNA synthesis or stability (Figure 10A). Compared to cells expressing wt NS1, the cells that expressed mutant NS1 had reduced expression of DNA-binding transcription factors, notably Myod1, Rara (retinoic acid receptor alpha), and the androgen receptor (Ar). On the other hand, the exclusion of NS1 from the nucleus caused the upregulation of 544 genes predominantly related to the innate immune response and pro-inflammatory pathways (Figure 10B). The mutant protein failed to repress relevant inflammatory components, including the Nlrp3 inflammasome, its associated cytokine IL-18, and central modulators of the NF-κB pathway (NFkB1 and NFkB1a). In addition, the absence of nuclear NS1 led to upregulation of Zc3h12a, which encodes the endoribonuclease MCPIP1 or Regnase-1. MCPIP1 is a critical immune homeostatic enzyme responsible for the degradation of both viral RNA and pro-inflammatory cytokine mRNAs (42–44). Furthermore, the downregulated fraction in the absence of nuclear NS1 was also enriched for terms associated with mitochondrial energy metabolism (Figure 10C). The top downregulated genes in the mitochondrial fraction included Pink1, encoding a key kinase required for mitophagy initiation, and genes encoding crucial subunits of the NADH dehydrogenase complex I (ND1, ND2, ND4, and ND6). This failure in genes related to mitochondrial quality control and oxidative phosphorylation, a known characteristic of severe dengue pathogenesis (45–47) led to a compensatory upregulation of the antioxidant Sod2 (superoxide dismutase), likely as the cell attempted to mitigate the resulting reactive oxygen species from dysfunctional mitochondria. This metabolic distress was accompanied by severe lipid dysregulation. The top upregulated genes in the lipid metabolism category were Pnpla2, encoding a key lipase driving lipolysis, and Cd36, encoding a fatty acid translocase. These results indicate that NS1 has a key function as a transcriptional regulator of the host cells to suppress antiviral inflammatory responses and promote a metabolic state favorable for viral replication.

**Figure 10.**
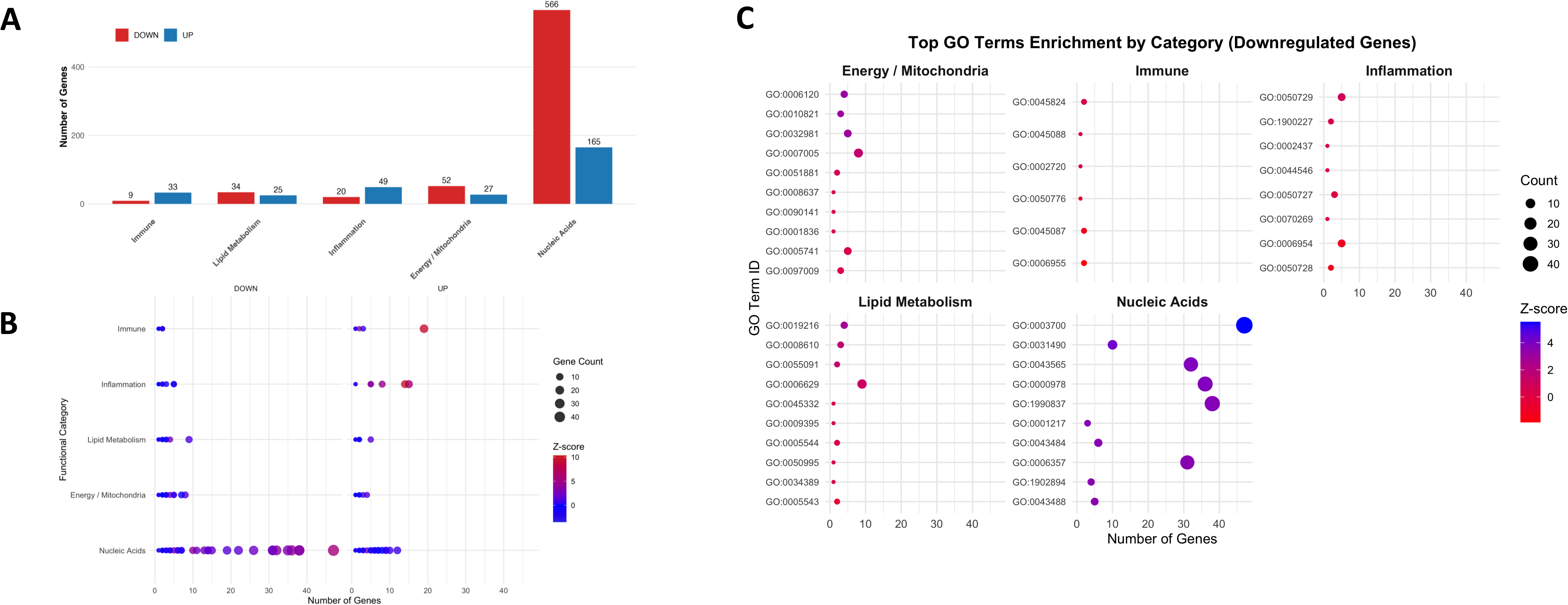
Gene ontology (GO) enrichment analysis of differentially expressed genes upon in BHK-21 cells expressing DENV2 wt-NS1 versus Mut3-NS1. **A**) Comparative GO category enrichment of upregulated and downregulated genes. The bar plot shows the total number of genes associated with each functional category for upregulated (blue) and downregulated (red) gene sets. The analysis reveals a predominant enrichment of *Nucleic Acid–related* categories compared with *Immune and Inflammatory processes*, as well as pathways associated with *Metabolism*, *Lipid Processing*, and *Mitochondrial Energy* functions. **B)** Dot plots showing functional category enrichment for upregulated and downregulated genes. The y-axis represents GO functional categories, and the x-axis indicates the number of genes associated with each term. Bubble size corresponds to gene count, and color represents the Z-score, reflecting relative enrichment strength. Upregulated (left facet) and downregulated (right facet) gene sets are shown separately, highlighting differences in functional enrichment across major biological processes, including *Nucleic Acid–related* categories; immune response, inflammation, metabolism, lipid processing, and mitochondrial energy pathways. **C)** Dot plot showing the top 10 enriched GO terms within each functional category (immune, inflammation, lipid metabolism, energy/mitochondria, nucleic acids) for downregulated genes. Each dot represents a GO term, with size proportional to the number of associated genes and color indicating the Z-score of enrichment, reflecting the magnitude of deviation from the expected gene count. Facets allow visualization of functional categories separately, highlighting category-specific enrichment patterns and enabling direct comparison of the most significantly affected biological processes.

## Discussion

A dual-localization model has traditionally defined the NS1 protein of orthoflaviviruses. Seminal studies established that intracellular NS1 resides predominantly within the lumen of the ER, as a dimer necessary for anchoring the viral replication complex to the ER membrane (18,48). In addition, a fraction of the mature NS1 glycoprotein is secreted into the extracellular space in association with lipids (3,8,49,50), and may be found in the sera of dengue patients at high concentration during the acute phase of disease. Under this classical view, the intracellular activity of NS1 was strictly limited to the ER and possibly also the cytoplasm (7). However, the findings presented in this study point to revising this view and adding NS1 to the list of DENV proteins that travel to the cell nucleus during infection, alongside NS3 and NS5.

Previous studies have reported circumstantial evidence of DENV NS1 in the nuclei of human endothelial (24) and mosquito cells (22,23). However, this study is the first to systematically validate these observations in a productive mammalian cell infection model. Notably, the time-course analysis demonstrates that the nuclear accumulation of DENV2 NS1 is a regulated early event that peaks concurrently with reported exponential viral RNA synthesis (51), effectively excluding the possibility that nuclear NS1 may be an artifact of apoptotic nuclear envelope degradation. The physical association of NS1 with the nuclear lamina (Lamin A/C and B1) and chromatin aligns with recent studies that identified nuclear pore complexes and epigenetic regulators as part of the NS1 interactome in both human and mosquito cells (25,26).

The NS1 dimer is about 100 kDa, exceeding the limits for passive diffusion through the nuclear pore complex (NPC). Indeed, pharmacological blockade with ivermectin, which is a recognized inhibitor of the importin α/β pathway (40,41,52), confirmed active nuclear entry of NS1. In addition, site-directed mutagenesis of conserved and surface-exposed NLS elements revealed that NS1 relies on the nuclear import machinery, similar to the DENV NS5 polymerase (53–55). The nuclear import machinery is also used by the ZIKV NS3 protein to reach the nuclear compartment (56). The mechanism by which NS1, located in the ER lumen, localizes to the cytoplasm to interact with the importin α/β machinery remains unclear; however, at least two different pathways could be proposed. First, exogenous NS1 may reach the cytosol via endosomal escape after internalization (14). Second, the endogenous pool of NS1 may undergo regulated retrotranslocation from the ER lumen to the cytosol, a mechanism previously described for other viral proteins, such as Rem-SP proteins from mouse mammary tumor virus, and toxins, such as CTA1 from *Vibrio cholerae* (57). In addition, the NS1 interaction with nuclear lamin A/C and B1 suggests that the translocation of NS1 may also occur via lateral displacement through the peripheral channels of the NPC. This mechanism allows integral membrane proteins to access the inner nuclear membrane without separating from the lipid bilayer, and it has been well-characterized for other high-molecular mass complexes requiring active, karyopherin-mediated transport (58). Remarkably, the absolute failure of the DENV2 harboring mutations in the NLS to replicate, and rescue of viability via trans-complementation, provides clear genetic evidence that the nuclear translocation of NS1 is not an accessory or artifactual phenomenon, but an obligate requirement for a productive viral life cycle. Indeed, mutation of position 189, which is part of the NLS of DENV NS1, has been reported to severely affect viral replication (4). An interesting observation is that the abortive infection obtained with the mutant virus, upon co-transfection with a plasmid expressing mutant NS1, indicates that although viral replication can be launched, the cell can contain the infection and prevent infectious virus release if NS1 fails to reach the nucleus.

The transcriptomic (RNA-seq) profiling of cells expressing the cytoplasm-restricted NS1 mutant and the wt NS1 revealed that nuclear NS1 functions as a transcriptional modulator. The functional category with the highest density of alterations was nucleic acid metabolism, indicating a profound reprogramming of the host’s transcriptional machinery. This demonstrates that the nuclear fraction of DENV2 NS1 does not merely shut down host transcription but also selectively upregulates specific host factors to its advantage. Interestingly, because androgen signaling acts as a negative regulator of the NF-κB pathway (59,60), the NS1-driven induction of Ar likely serves as a critical, hijacked molecular brake to suppress host inflammation. Moreover, several genes repressed by the wt NS1 are upregulated when the mutant NS1 is present, indicating a certain reciprocal regulatory interplay in the genes activated or repressed by the wt NS1 versus the mutated NS1. For example, by actively suppressing Zc3h12a transcription during a DENV infection, nuclear NS1 may secure a dual advantage: protection of the viral genome from enzymatic degradation, while simultaneously preventing the resolution of the inflammatory response, upregulated by the mutant NS1.

The role of nuclear NS1 appears to vary with the intracellular location and cell type. Exposure of human monocytes to DENV2 wt NS1 at high concentrations (10µg/mL) actively induced pro-inflammatory and antiviral transcriptomic responses (61). In this study, a pro-inflammatory response was mainly observed with the cytoplasm-restricted mutant, while the anti-viral mainly observed with the wt NS1. Increased cytokine expression has been attributed to extracellular NS1 signaling via cell-surface receptors such as TLR4 (11). Our results show that exogenous NS1 is efficiently trafficked to the nucleus. Consequently, the cytokine dysregulation seen in severe dengue may not depend exclusively on transmembrane signaling cascades but rather on direct, active transcriptional reprogramming executed by internalized NS1 within the nucleus. Besides, exposure of cultured cells to soluble NS1 renders them more susceptible to DENV infection (14,15), yet the reasons for this remain poorly understood. Our data, indicating that soluble NS1 can reach the cell nucleus and modulate gene expression to favor viral replication, provide at least a partial mechanistic explanation for this phenomenon.

On the other hand, the GO frequency analysis highlighted severe transcriptomic alterations in the energy/mitochondria and lipid metabolism categories. The extranuclear mutant NS1 appears to cause a profound metabolic collapse and lipid deregulation. Orthoflavivirus replication relies heavily on the continuous recruitment of lipids for the biogenesis of replication organelles and viral envelopes (62,63). The exacerbated lipolytic profile (Pnpla2) and increased lipid uptake markers (Cd36) in the cytoplasm-restricted mutant indicate a decompensated metabolic stress state rather than efficient recruitment of resources. Again, these results are compatible with a model of spatial metabolic cooperation: while cytoplasmic NS1 interacts with enzymes like GAPDH to accelerate glycolysis and provide rapid biosynthetic precursors (64), nuclear NS1 transcriptionally stabilizes oxidative phosphorylation and prevents lipotoxic stress. This bipartite regulation would ensure that the host cell meets the massive energetic demands of viral replication without succumbing to premature oxidative stress-induced apoptosis.

While this study offers a comprehensive characterization of nuclear activities of NS1, it is limited to DENV2 and the highly permissive BHK-21 cell line. Thus, future studies must validate these findings with other orthoflaviviruses and cell types. However, preliminary evidence obtained from Huh-7 cells infected with DENV2 and BHK-21 cells infected with DENV4 indicates that NS1 also reaches the nucleus in those infections (data not shown). Also, mechanistic validation of specific transcriptional targets, such as MCPIP1 and the mitochondrial subunits, in primary human target cells is needed. Nonetheless, establishing the absolute dependence of viral viability on the nuclear import of NS1 exposes the NLS-importin interface as a highly specific and vulnerable therapeutic target for dengue infection. Furthermore, the rational attenuation of DENV via targeted mutations in the NS1 NLS offers a promising strategy for developing next-generation live-attenuated vaccines.

## Acknowledgments

We like to acknowledge Dr. Ana Lorena Gutiérrez Escolano and Ana C. Alcalá for their critical reading of the manuscript and M.Sc. Alan Perez Hernandez for his support in obtaining the confocal microscope images.

## References

1. Gubler DJ. Dengue and dengue hemorrhagic fever. Clin Microbiol Rev. 1998 Jul;11(3):480–96. doi:10.1128/CMR.11.3.480

2. Lai NS. Roles and prospects of dengue virus nonstructural proteins as antiviral targets: an easy digest. MJMS. 2018;25(5):6–15. doi:10.21315/mjms2018.25.5.2

3. Akey DL, Brown WC, Dutta S, Konwerski J, Jose J, Jurkiw TJ, et al. Flavivirus NS1 structures reveal surfaces for associations with membranes and the immune system. Science. 2014 Feb 21;343(6173):881–5. doi:10.1126/science.1247749

4. Winkelmann ER, Widman DG, Suzuki R, Mason PW. Analyses of mutations selected by passaging a chimeric flavivirus identify mutations that alter infectivity and reveal an interaction between the structural proteins and the nonstructural glycoprotein NS1. Virology. 2011 Dec;421(2):96–104. doi:10.1016/j.virol.2011.09.007

5. Scaturro P, Cortese M, Chatel-Chaix L, Fischl W, Bartenschlager R. Dengue virus non-structural protein 1 modulates infectious particle production via interaction with the structural proteins. PLoS Pathog. 2015 Nov 12;11(11):e1005277. doi:10.1371/journal.ppat.1005277

6. Jacobs MG, Robinson PJ, Bletchly C, Mackenzie JM, Young PR. Dengue virus nonstructural protein 1 is expressed in a glycosyl-phosphatidylinositol-linked form that is capable of signal transduction. FASEB J. 2000 Aug;14(11):1603–10. doi:10.1096/fj.99-0829com

7. Castillo JM, Cruz-Pérez R, Talamás-Lara D, Ludert JE. Kinesin light chain 1 interacts with NS1 and is a susceptibility factor for dengue virus infection in mosquito cells. J Gen Virol. 2025 Jul 16;106(7). doi:10.1099/jgv.0.002132

8. Gutsche I, Coulibaly F, Voss JE, Salmon J, d’Alayer J, Ermonval M, et al. Secreted dengue virus nonstructural protein NS1 is an atypical barrel-shaped high-density lipoprotein. Proc Natl Acad Sci USA. 2011 May 10;108(19):8003–8. doi:10.1073/pnas.1017338108

9. Flamand M, Megret F, Mathieu M, Lepault J, Rey FA, Deubel V. Dengue virus type 1 nonstructural glycoprotein NS1 is secreted from mammalian cells as a soluble hexamer in a glycosylation-dependent fashion. J Virol. 1999 Jul;73(7):6104–10. doi:10.1128/JVI.73.7.6104-6110.1999

10. Alcon S, Talarmin A, Debruyne M, Falconar A, Deubel V, Flamand M. Enzyme-linked immunosorbent assay specific to dengue virus type 1 nonstructural protein NS1 reveals circulation of the antigen in the blood during the acute phase of disease in patients experiencing primary or secondary infections. J Clin Microbiol. 2002 Feb;40(2):376–81. doi:10.1128/JCM.40.02.376-381.2002

11. Modhiran N, Watterson D, Muller DA, Panetta AK, Sester DP, Liu L, et al. Dengue virus NS1 protein activates cells via Toll-like receptor 4 and disrupts endothelial cell monolayer integrity. Sci Transl Med. 2015 Sep 9;7(304). doi:10.1126/scitranslmed.aaa3863

12. Avirutnan P, Fuchs A, Hauhart RE, Somnuke P, Youn S, Diamond MS, et al. Antagonism of the complement component C4 by flavivirus nonstructural protein NS1. J Exp Med. 2010 Apr 12;207(4):793–806. doi:10.1084/jem.20092545

13. Beatty PR, Puerta-Guardo H, Killingbeck SS, Glasner DR, Hopkins K, Harris E. Dengue virus NS1 triggers endothelial permeability and vascular leak that is prevented by NS1 vaccination. Sci Transl Med. 2015 Sep 9;7(304). doi:10.1126/scitranslmed.aaa3787

14. Alcon-LePoder S, Drouet MT, Roux P, Frenkiel MP, Arborio M, Durand-Schneider AM, et al. The secreted form of dengue virus nonstructural protein NS1 is endocytosed by hepatocytes and accumulates in late endosomes: implications for viral infectivity. J Virol. 2005 Sep;79(17):11403–11. doi:10.1128/JVI.79.17.11403-11411.2005

15. Alayli F, Scholle F. Dengue virus NS1 enhances viral replication and pro-inflammatory cytokine production in human dendritic cells. Virology. 2016 Sep;496:227–36. doi:10.1016/j.virol.2016.06.008

16. Falconar AKI. The dengue virus nonstructural-1 protein (NS1) generates antibodies to common epitopes on human blood clotting, integrin/adhesin proteins and binds to human endothelial cells: potential implications in haemorrhagic fever pathogenesis. Arch Virol. 1997 May;142(5):897–916. doi:10.1007/s007050050127

17. Lin C, Lei H, Shiau A, Liu C, Liu H, Yeh T, et al. Antibodies from dengue patient sera cross-react with endothelial cells and induce damage. J Med Virol. 2003 Jan;69(1):82–90. doi:10.1002/jmv.10261

18. Muller DA, Young PR. The flavivirus NS1 protein: Molecular and structural biology, immunology, role in pathogenesis and application as a diagnostic biomarker. Antiviral Res. 2013 May;98(2):192–208. doi:10.1016/j.antiviral.2013.03.008

19. Chen HR, Lai YC, Yeh TM. Dengue virus non-structural protein 1: a pathogenic factor, therapeutic target, and vaccine candidate. J Biomed Sci. 2018 Dec;25(1):58. doi:10.1186/s12929-018-0462-0

20. Rastogi M, Sharma N, Singh SK. Flavivirus NS1: a multifaceted enigmatic viral protein. Virol J. 2016 Dec;13(1):131. doi:10.1186/s12985-016-0590-7

21. Alcalá AC, Ludert JE. The dengue virus NS1 protein; new roles in pathogenesis due to similarities with and affinity for the high-density lipoprotein (HDL)?. PLoS Pathog. 2023 Aug 24;19(8):e1011587. doi:10.1371/journal.ppat.1011587

22. Reyes-Ruiz JM, Osuna-Ramos JF, Cervantes-Salazar M, Lagunes Guillen AE, Chávez-Munguía B, Salas-Benito JS, et al. Strand-like structures and the nonstructural proteins 5, 3 and 1 are present in the nucleus of mosquito cells infected with dengue virus. Virology. 2018 Feb;515:74–80. doi:10.1016/j.virol.2017.12.014

23. Alcalá AC, Medina F, González-Robles A, Salazar-Villatoro L, Fragoso-Soriano RJ, Vásquez C, et al. The dengue virus non-structural protein 1 (NS1) is secreted efficiently from infected mosquito cells. Virology. 2016 Jan;488:278–87. doi:10.1016/j.virol.2015.11.020

24. Poungsawai J, Kanlaya R, Pattanakitsakul S-Nga, Thongboonkerd V. Subcellular localizations and time-course expression of dengue envelope and non-structural 1 proteins in human endothelial cells. Microb Pathog. 2011 Sep;51(3):225–9. doi:10.1016/j.micpath.2011.04.011

25. Caraballo GI, Rosales R, Viettri M, Castillo JM, Cruz R, Ding S, et al. The dengue virus nonstructural protein 1 (NS1) interacts with the putative epigenetic regulator DIDO1 to promote flavivirus replication in mosquito cells. J Virol. 2022 Jun 22;96(12):e00704–22. doi:10.1128/jvi.00704-22

26. Hafirassou ML, Meertens L, Umaña-Diaz C, Labeau A, Dejarnac O, Bonnet-Madin L, et al. A global interactome map of the dengue virus NS1 identifies virus restriction and dependency host factors. Cell Reports. 2017 Dec;21(13):3900–13. doi:10.1016/j.celrep.2017.11.094

27. Sawasdichai A, Chen HT, Abdul Hamid N, Jayaraman PS, Gaston K. In situ subcellular fractionation of adherent and non-adherent mammalian cells. JoVE. 2010 Jul 23;(41):1958. doi:10.3791/1958

28. Arndt C, Koristka S, Feldmann A, Bachmann M. Native Polyacrylamide Gels. In: Kurien BT, Scofield RH, editors. Electrophoretic separation of proteins [Internet]. New York, NY: Springer New York; 2019 [cited 2025 Nov 28]. p. 87–91. (Methods in Molecular Biology). Available from: http://link.springer.com/10.1007/978-1-4939-8793-1_8 doi:10.1007/978-1-4939-8793-1_8

29. Kosugi S, Hasebe M, Tomita M, Yanagawa H. Systematic identification of cell cycle-dependent yeast nucleocytoplasmic shuttling proteins by prediction of composite motifs. Proc Natl Acad Sci USA. 2009 Jun 23;106(25):10171–6. doi:10.1073/pnas.0900604106

30. Waterhouse AM, Procter JB, Martin DMA, Clamp M, Barton GJ. Jalview Version 2—a multiple sequence alignment editor and analysis workbench. Bioinformatics. 2009 May 1;25(9):1189–91. doi:10.1093/bioinformatics/btp033

31. Jumper J, Evans R, Pritzel A, Green T, Figurnov M, Ronneberger O, et al. Highly accurate protein structure prediction with AlphaFold. Nature. 2021 Aug 26;596(7873):583–9. doi:10.1038/s41586-021-03819-2

32. Pettersen EF, Goddard TD, Huang CC, Couch GS, Greenblatt DM, Meng EC, et al. UCSF Chimera—A visualization system for exploratory research and analysis. J Comput Chem. 2004 Oct;25(13):1605–12. doi:10.1002/jcc.20084

33. Mikaelian I, Sergeant A. Modification of the overlap extension method for extensive mutagenesis on the same template. In: in vitro mutagenesis protocols [Internet]. New Jersey: Humana Press; 1996 [cited 2025 Nov 21]. p. 193–202. Available from: http://link.springer.com/10.1385/0-89603-332-5:193 doi:10.1385/0-89603-332-5:193

34. Cherkashchenko L, Gros N, Trausch A, Neyret A, Hénaut M, Dubois G, et al. Validation of flavivirus infectious clones carrying fluorescent markers for antiviral drug screening and replication studies. Front Microbiol. 2023 Sep 15;14:1201640. doi:10.3389/fmicb.2023.1201640

35. Kim D, Langmead B, Salzberg SL. HISAT: a fast spliced aligner with low memory requirements. Nat Methods. 2015 Apr;12(4):357–60. doi:10.1038/nmeth.3317

36. Yu G, Wang LG, Han Y, He QY. clusterProfiler: an R package for comparing biological themes among gene clusters. OMICS. 2012 May 1;16(5):284–7. doi:10.1089/omi.2011.0118

37. Durinck S, Spellman PT, Birney E, Huber W. Mapping identifiers for the integration of genomic datasets with the R/Bioconductor package biomaRt. Nat Protoc. 2009 Aug;4(8):1184–91. doi:10.1038/nprot.2009.97

38. Jovanovic-Talisman T, Zilman A. Protein transport by the nuclear pore complex: Simple biophysics of a complex biomachine. Biophys J. 2017 Jul;113(1):6–14. doi:10.1016/j.bpj.2017.05.024

39. Mohr D, Frey S, Fischer T, Güttler T, Görlich D. Characterisation of the passive permeability barrier of nuclear pore complexes. EMBO J. 2009 Sep 2;28(17):2541–53. doi:10.1038/emboj.2009.200

40. Mastrangelo E, Pezzullo M, De Burghgraeve T, Kaptein S, Pastorino B, Dallmeier K, et al. Ivermectin is a potent inhibitor of flavivirus replication specifically targeting NS3 helicase activity: new prospects for an old drug. J Antimicrob Chemother. 2012 Aug 1;67(8):1884–94. doi:10.1093/jac/dks147

41. Wagstaff KM, Sivakumaran H, Heaton SM, Harrich D, Jans DA. Ivermectin is a specific inhibitor of importin α/β-mediated nuclear import able to inhibit replication of HIV-1 and dengue virus. Biochem J. 2012 May 1;443(3):851–6. doi:10.1042/BJ20120150

42. Akira S. Regnase-1, a ribonuclease involved in the regulation of immune responses. Cold Spring Harb Symp Quant Biol. 2013 Jan 1;78(0):51–60. doi:10.1101/sqb.2013.78.019877

43. Mino T, Murakawa Y, Fukao A, Vandenbon A, Wessels HH, Ori D, et al. Regnase-1 and roquin regulate a common element in inflammatory mRNAs by spatiotemporally distinct mechanisms. Cell. 2015 May;161(5):1058–73. doi:10.1016/j.cell.2015.04.029

44. Mao R, Yang R, Chen X, Harhaj EW, Wang X, Fan Y. Regnase-1, a rapid response ribonuclease regulating inflammation and stress responses. Cell Mol Immunol. 2017 May;14(5):412–22. doi:10.1038/cmi.2016.70

45. Singh B, Avula K, Sufi SA, Parwin N, Das S, Alam MF, et al. Defective mitochondrial quality control during dengue infection contributes to disease pathogenesis. J Virol. 2022 Oct 26;96(20):e00828–22. doi:10.1128/jvi.00828-22

46. Chatel-Chaix L, Cortese M, Romero-Brey I, Bender S, Neufeldt CJ, Fischl W, et al. Dengue virus perturbs mitochondrial morphodynamics to dampen innate immune responses. Cell Host & Microbe. 2016 Sep;20(3):342–56. doi:10.1016/j.chom.2016.07.008

47. Ahmed S, Varga RD, Yang J. The impacts of dengue virus infection on mitochondrial functions and dynamics. IJMS. 2025 Sep 15;26(18):8968. doi:10.3390/ijms26188968

48. Winkler G, Randolph VB, Cleaves GR, Ryan TE, Stollar V. Evidence that the mature form of the flavivirus nonstructural protein NS1 is a dimer. Virology. 1988 Jan;162(1):187–96. doi:10.1016/0042-6822(88)90408-4

49. Benfrid S, Park K, Dellarole M, Voss JE, Tamietti C, Pehau-Arnaudet G, et al. Dengue virus NS1 protein conveys pro-inflammatory signals by docking onto high-density lipoproteins. EMBO Reports. 2022 Jul 5;23(7):e53600. doi:10.15252/embr.202153600

50. Chew BLA, Ngoh AQ, Phoo WW, Chan KWK, Ser Z, Tulsian NK, et al. Secreted dengue virus NS1 from infection is predominantly dimeric and in complex with high-density lipoprotein. eLife. 2024 May 24;12:RP90762. doi:10.7554/eLife.90762

51. Neufeldt CJ, Cortese M, Acosta EG, Bartenschlager R. Rewiring cellular networks by members of the Flaviviridae family. Nat Rev Microbiol. 2018 Mar;16(3):125–42. doi:10.1038/nrmicro.2017.170

52. Yang SNY, Atkinson SC, Wang C, Lee A, Bogoyevitch MA, Borg NA, et al. The broad spectrum antiviral ivermectin targets the host nuclear transport importin α/β1 heterodimer. Antiviral Res. 2020 May;177:104760. doi:10.1016/j.antiviral.2020.104760

53. Rawlinson SM, Pryor MJ, Wright PJ, Jans DA. CRM1-mediated nuclear export of dengue virus RNA polymerase NS5 modulates interleukin-8 induction and virus production. J Biol Chem. 2009 Jun;284(23):15589–97. doi:10.1074/jbc.M808271200

54. Pryor MJ, Rawlinson SM, Butcher RE, Barton CL, Waterhouse TA, Vasudevan SG, et al. Nuclear localization of dengue virus nonstructural protein 5 through its importin α/β–recognized nuclear localization sequences is integral to viral infection. Traffic. 2007 Jul;8(7):795–807. doi:10.1111/j.1600-0854.2007.00579.x

55. Tay MYF, Fraser JE, Chan WKK, Moreland NJ, Rathore AP, Wang C, et al. Nuclear localization of dengue virus (DENV) 1–4 non-structural protein 5; protection against all 4 DENV serotypes by the inhibitor Ivermectin. Antiviral Res. 2013 Sep;99(3):301–6. doi:10.1016/j.antiviral.2013.06.002

56. De Jesús-González LA, Palacios-Rápalo SN, Reyes-Ruiz JM, Osuna-Ramos JF, Farfán-Morales CN, Cordero-Rivera CD, et al. Nucleo-cytoplasmic transport of ZIKV non-structural 3 protein is mediated by importin-α/β and exportin CRM-1. J Virol. 2023 Jan 31;97(1):e01773–22. doi:10.1128/jvi.01773-22

57. Byun H, Gou Y, Zook A, Lozano MM, Dudley JP. ERAD and how viruses exploit it. Front Microbiol. 2014 Jul 3;5. doi:10.3389/fmicb.2014.00330

58. King MC, Lusk C, Blobel G. Karyopherin-mediated import of integral inner nuclear membrane proteins. Nature. 2006 Aug;442(7106):1003–7. doi:10.1038/nature05075

59. Basílio J, Hochreiter B, Hoesel B, Sheshori E, Mussbacher M, Hanel R, et al. Antagonistic functions of androgen receptor and NF-κB in prostate cancer—experimental and computational analyses. Cancers. 2022 Dec 14;14(24):6164. doi:10.3390/cancers14246164

60. Zhang L, Altuwaijri S, Deng F, Chen L, Lal P, Bhanot UK, et al. NF-κB regulates androgen receptor expression and prostate cancer growth. Am J Pathol. 2009 Aug;175(2):489–99. doi:10.2353/ajpath.2009.080727

61. Saisingha K, Jearanaiwitayakul T, Watterson D, Modhiran N, Ponpuak M, Ubol S. Transcriptome analysis of monocytes treated with dengue virus nonstructural protein 1 revealed a shift in transcripts involved in self-propagated proinflammation and antiviral responses. J Infect Dis. 2025 Jul 11;231(6):e1170–82. doi:10.1093/infdis/jiaf166

62. Heaton NS, Randall G. Dengue virus-induced autophagy regulates lipid metabolism. Cell Host & Microbe. 2010 Nov;8(5):422–32. doi:10.1016/j.chom.2010.10.006

63. Perera R, Riley C, Isaac G, Hopf-Jannasch AS, Moore RJ, Weitz KW, et al. Dengue virus infection perturbs lipid homeostasis in infected mosquito cells. PLoS Pathog. 2012 Mar 22;8(3):e1002584. doi:10.1371/journal.ppat.1002584

64. Allonso D, Andrade IS, Conde JN, Coelho DR, Rocha DCP, Da Silva ML, et al. Dengue virus NS1 protein modulates cellular energy metabolism by increasing glyceraldehyde-3-phosphate dehydrogenase activity. J Virol. 2015 Dec;89(23):11871–83. doi:10.1128/JVI.01342-15

