## Supplemental Figures for "Dengue virus NS1 undergoes partial nuclear translocation to modulate host transcription and support viral replication"

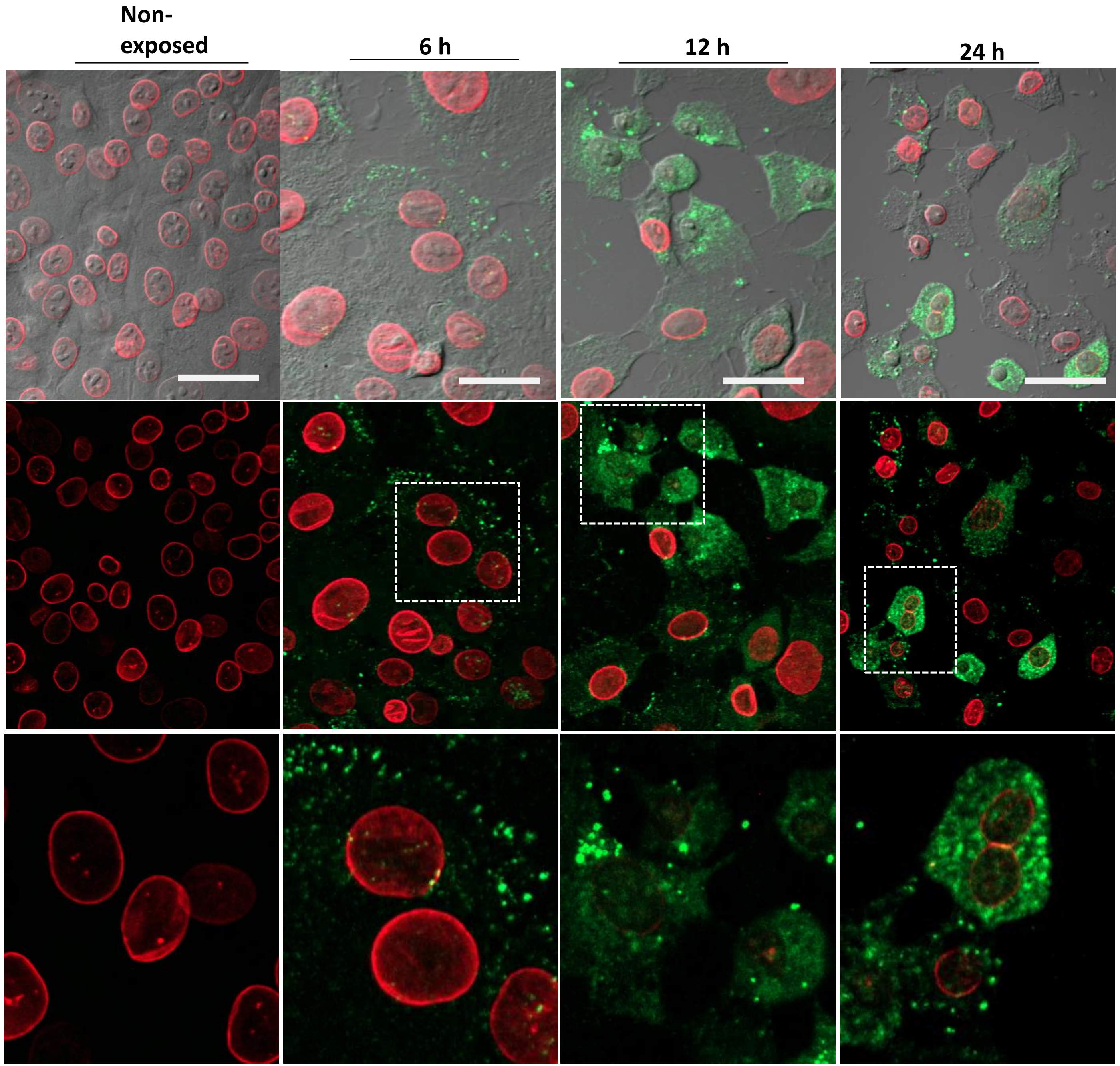


**Supplemental Figure 1. Internalization of recombinant NS1 in BHK-21 cells.** BHK-21 cells monolayers were incubated with soluble, recombinant DENV-2 NS1 protein (3 µg/ml) for 6, 12, and 24 h, then fixed and processed for confocal microscopy. NS1 protein was visualized using an anti-NS1 Mab and an Alexa 488 (green), and nuclear lamina B1 using a commercial monoclonal antibody and Alexa 647 (red), as primary and secondary antibodies. The dotted area in the middle panels is enlarged in the bottom panels. Scale bar: 10 µm

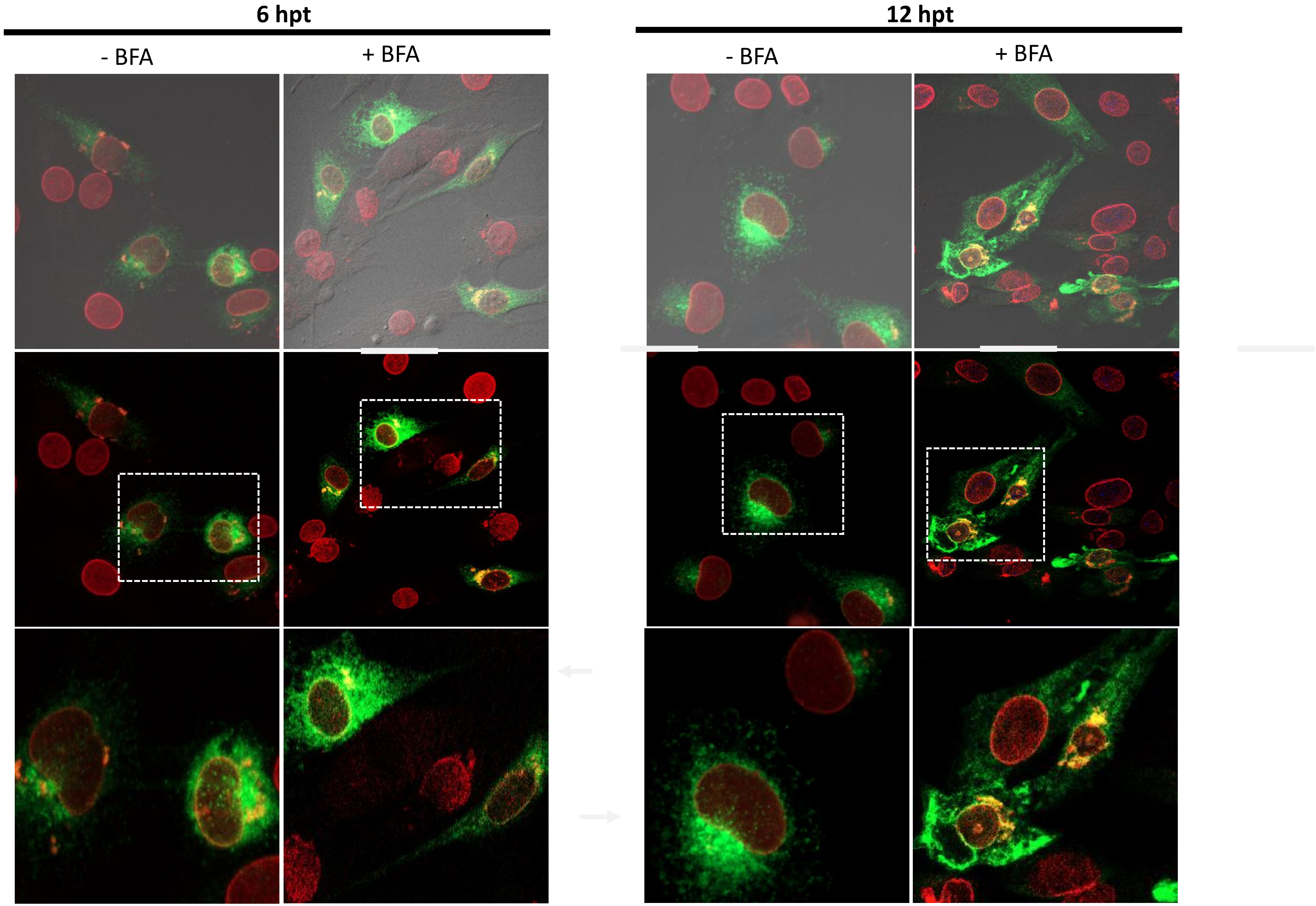


**Supplemental Figure 2. Nuclear trafficking of NS1 in brefeldin A-treated cells.** BHK-21 cells transfected with plasmid encoding wt NS1 were treated or not with Brefeldin A (3 µg/ml) and at the indicated times, fixed and processed for confocal microscopy. NS1 protein was visualized using an anti-NS1 Mab and an Alexa 488 (green), and nuclear lamina B1 using a commercial monoclonal antibody and Alexa 647 (red), as primary and secondary antibodies. Dotted area in the middle panels enlarged in the bottom panels. Scale bar: 10 µm.

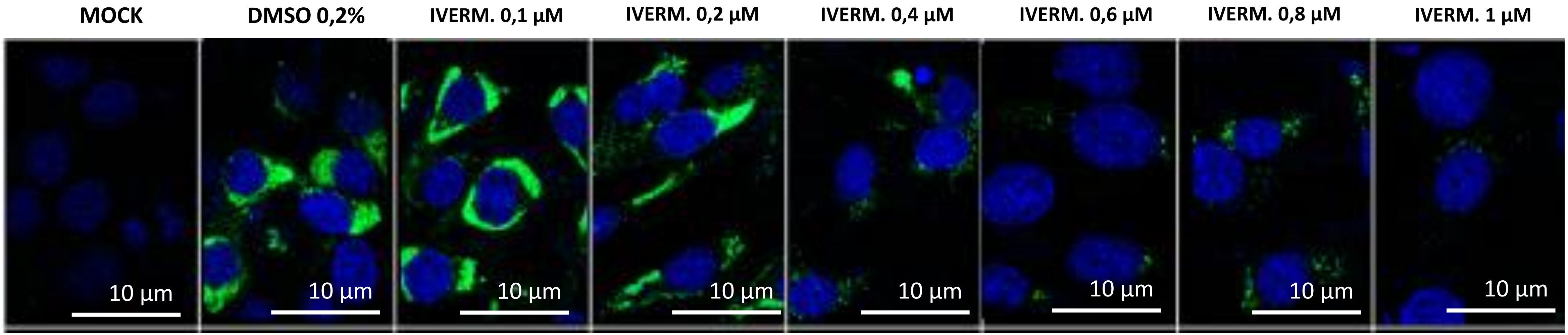


**Supplemental Figure 3. Dose-dependent reduction of DENV2 NS1 protein expression by Ivermectin treatment**. Immunofluorescence microscopy images of BHK-21 cells, mock infected or infected with DENV2 (MOI=3), treated with ivermectin at different concentrations as indicated or not (DMSO 0,2%), and fixed 24 hours post-infection (hpi). Cell nuclei were stained with DAPI (blue), and viral NS1 protein was detected using a mouse anti-NS1 Mab as primary antibody and an Alexa 488 fluorophore (green) conjugated anti-mouse Mab. Scale bar: 10 µm for all panels.

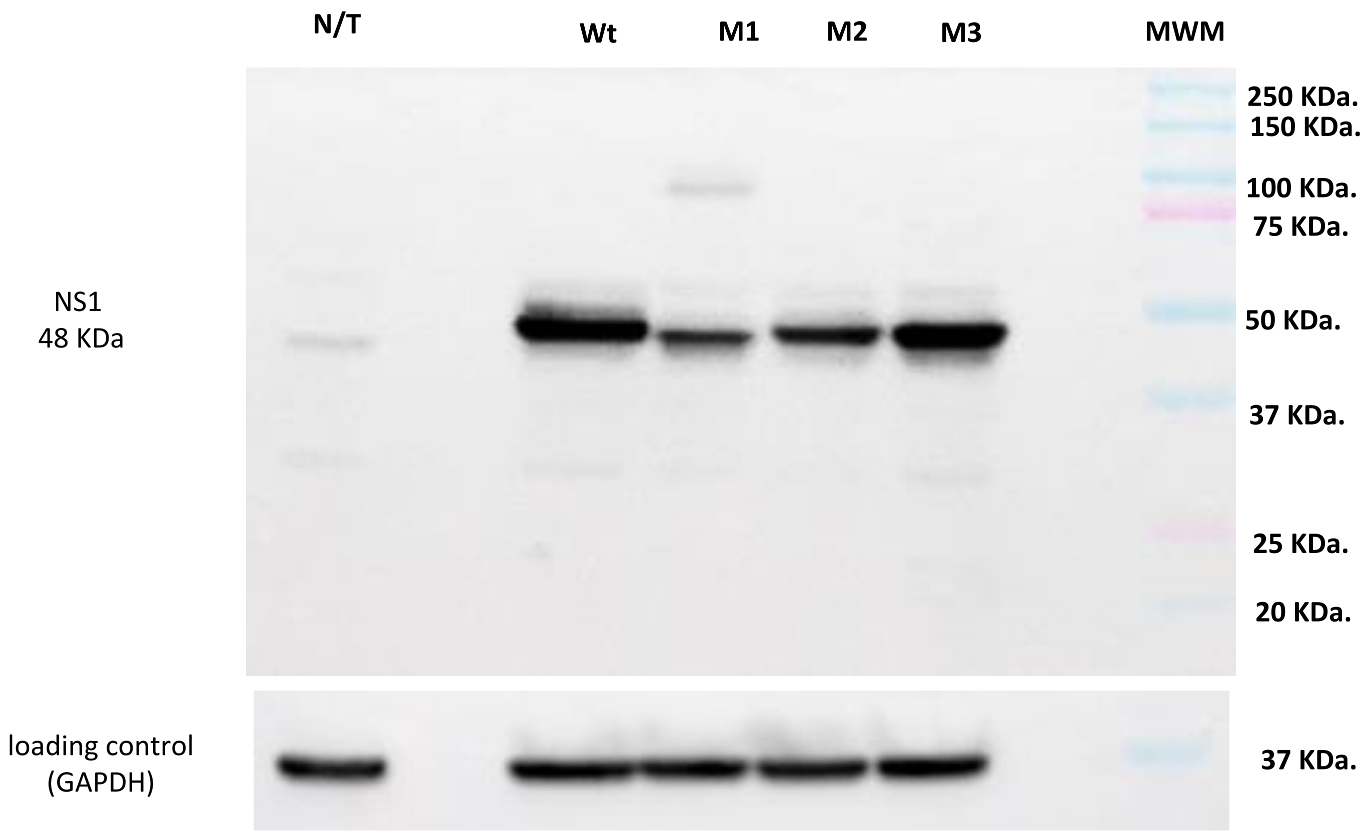


**Supplemental Figure 4. Expression of recombinant wild-type, and mutants M1, M2, and M3 NS1.** BHK-21 cells were transfected with plasmids encoding Wild-type (Wt) or mutants (M1, M2, and M3) NS1, harvested at 24 hours post-transfection, and total protein extracts (20 µg) were analyzed by Western blot. GAPDH (37 kDa) was used as a loading control. N/T: non-transfected cells; MWM: molecular protein markers. No significant differences in expression levels were observed between wt-NS1 and the Mut3-NS1 variants.

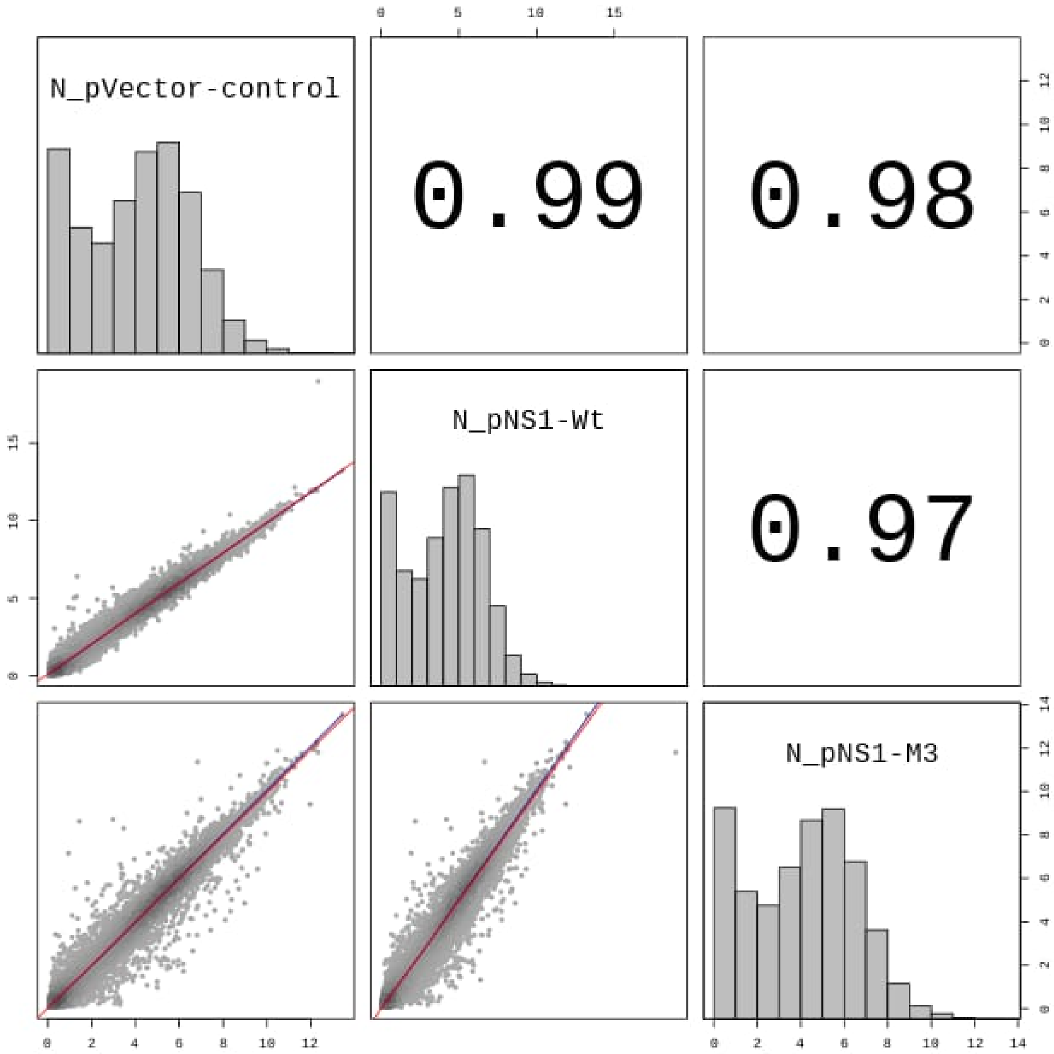


**Supplemental Figure 5. Correlation analysis of global gene expression between NS1 variants and the control.** Pairwise scatter plot comparing the transcriptomic profiles of cells transfected with the empty vector (pVector-control), wild-type NS1 (pNS1-Wt), and the Mut3-NS1 mutant (pNS1-M3). The diagonal shows the expression distribution histograms for each condition. The upper right panel indicates the Pearson correlation coefficients (r) between the compared pairs, demonstrating high overall similarity (r > 0.97). The lower left panel shows the scatter plots of normalized gene expression levels, where each point represents an individual gene; the red line indicates the linear regression trend.

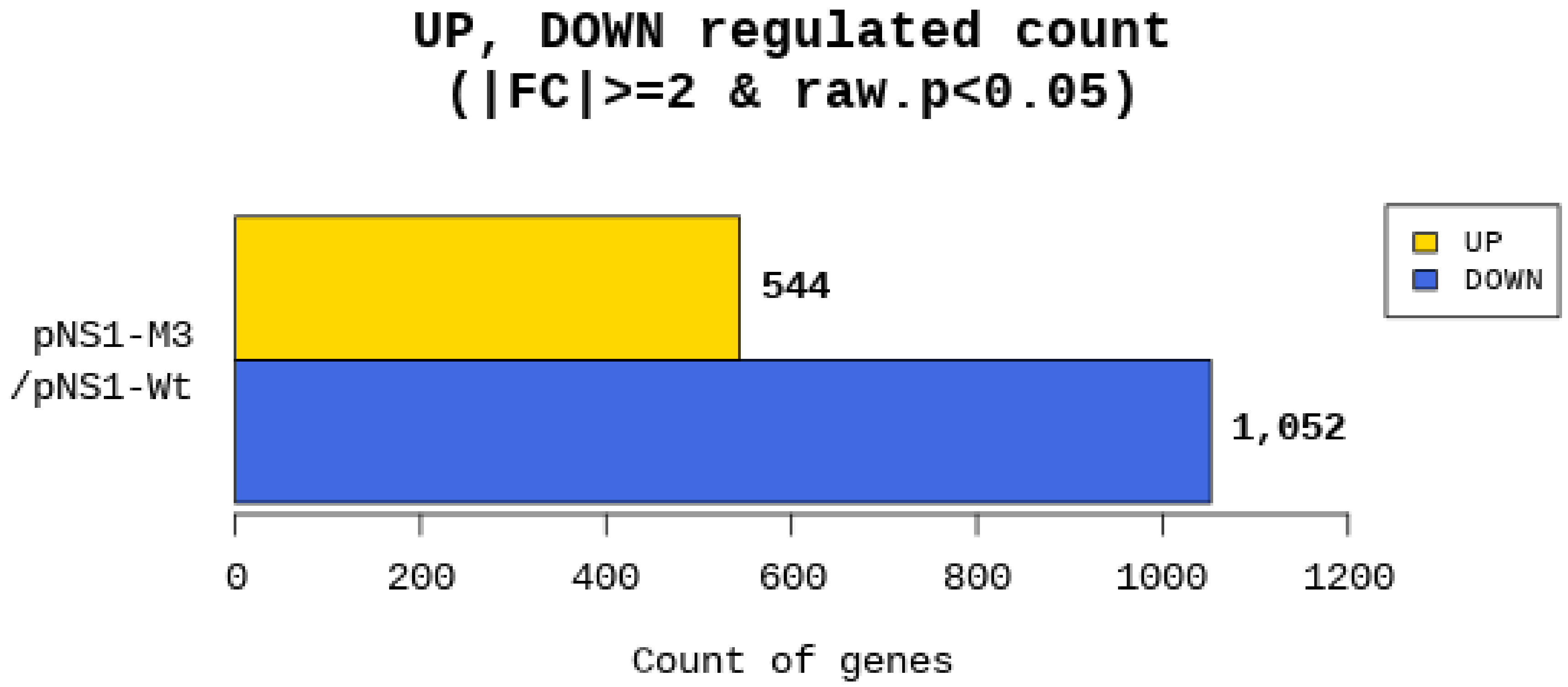


**Supplemental Figure 6**. **Bar chart showing the total number of significantly upregulated and downregulated differentially expressed genes (DEGs) in BHK-21 cells expressing Mut3-NS1 (NS1-M3) and wt-NS1 (NS1-Wt).**
