## Supplemental Table for "Dengue virus NS1 undergoes partial nuclear translocation to modulate host transcription and support viral replication"

**SUPPLEMENTAL TABLE.** Primers used in this study to detect (NS1), mutate (MUT) and sequence (SECNS1) the DENV2 NS1 gene.

| PRIMER | SECUENCES |
|---|---|
| NS1F | 5'-AGAATTCGAGCTCACCATGGGATC-3' |
| NS1R | 5'-CTGCTAGCTCGAGTCAGGCATAATC-3' |
| MUT1F | 5'-GCGGCCATACAGGACAACCAGGCCGTCCAT-3' |
| MUT1R | 5'-ATGGACGGCCTGGTTGTCCTGTATGGCCGC-3' |
| MUT2F | 5'-GACACATGGAAGATAGAGAAAGCCTCTTTC-3' |
| MUT2R | 5'-GAAAGAGGCTTTCTCTATCTTCCATGTGTC-3' |
| SECNS1F | 5'-GGCGGGTCTAGAGCCTCTGCT-3' |
| SECNS1R1 | 5'-CCAGAAGTCAGATGCTCAAGG-3' |
| SECNS1R2 | 5'-GTTGAGACACTGGTCCAGCG-3' |
